# Glucocorticoids suppress NF-κB-mediated neutrophil control of *Aspergillus fumigatus* hyphal growth

**DOI:** 10.1101/2023.12.21.572739

**Authors:** Savini U. Thrikawala, Molly Anderson, Emily E. Rosowski

**Affiliations:** Department of Biological Sciences, Clemson University, Clemson, South Carolina, United States of America

## Abstract

Glucocorticoids are a major class of therapeutic anti-inflammatory and immunosuppressive drugs prescribed to patients with inflammatory diseases, to avoid transplant rejection, and as part of cancer chemotherapy. However, exposure to these drugs increases the risk of opportunistic infections such as with the fungus *Aspergillus fumigatus*. Prolonged glucocorticoid therapy is one of the main risks for invasive aspergillosis, which causes mortality in >50% of infected patients. The mechanisms by which glucocorticoids increase susceptibility to *A. fumigatus* are poorly understood. Here, we used a zebrafish larva-*Aspergillus* infection model to identify innate immune mechanisms altered by glucocorticoid treatment. Infected larvae exposed to dexamethasone succumb to the infection at a significantly higher rate than control larvae. However, both macrophages and neutrophils are still recruited to the site of infection and dexamethasone treatment does not significantly affect fungal spore killing. Instead, the primary effect of dexamethasone manifests later in infection with treated larvae exhibiting increased invasive hyphal growth. In line with this, dexamethasone predominantly inhibits neutrophil function, rather than macrophage function. Dexamethasone-induced mortality also depends on the glucocorticoid receptor. One pathway that glucocorticoids can inhibit is NF-κB activation and we report that dexamethasone partially suppresses NF-κB activation at the infection site by inducing the transcription of IκB via the glucocorticoid receptor. Independent CRISPR/Cas9 targeting of IKKγ to prevent NF-κB activation also increases invasive *A. fumigatus* growth and larval mortality. However, dexamethasone treatment of IKKγ crispant larvae further increases invasive hyphal growth, suggesting that dexamethasone may suppress other pathways in addition to NF-κB to promote host susceptibility. Collectively, we find that dexamethasone acts through the glucocorticoid receptor to suppress NF-κB-mediated neutrophil control of *A. fumigatus* hyphae in zebrafish larvae.

**Author Summary:** Glucocorticoids are drugs that stop inflammation and suppress the immune system. Glucocorticoids are effective in treating inflammatory diseases such as asthma and arthritis, preventing organ rejection after transplant surgery, and in ameliorating the side effects of cancer chemotherapy. However, as these drugs suppress the immune system, patients taking glucocorticoids are more prone to infections such as with the environmental fungus *Aspergillus fumigatus*. The specific mechanisms that glucocorticoids inhibit to increase susceptibility to infection are largely unknown. Here, we used a larval zebrafish model of *A. fumigatus* infection to determine that glucocorticoids mainly suppress the ability of neutrophils to control the fungal hyphal growth that causes tissue damage. Our study provides insight into future strategies to treat *A. fumigatus* infection in patients undergoing glucocorticoid therapy.

## Introduction

Glucocorticoids are potent immunosuppressive and anti-inflammatory drugs that are prescribed for a range of conditions, including chronic inflammation, lymphoid malignancies, autoimmune conditions, and to avoid rejection in bone marrow and solid organ transplant patients [1–3]. However, prolonged use of glucocorticoids causes adverse effects such as metabolic disorders, hypertension, osteoporosis, and depression [4]. The immunosuppressive effects of glucocorticoids also increase the risk of opportunistic infections [4]. Invasive aspergillosis caused by *Aspergillus fumigatus* is the most common fungal infection associated with glucocorticoid therapy [5]. While immunocompetent hosts effectively clear inhaled airborne *A. fumigatus* spores from the lungs and airways, in patients undergoing glucocorticoid therapy spores can germinate into invasive filamentous hyphae, destroying tissues and organs [6]. Anti-fungal treatments are often ineffective, partially due to growing drug resistance among fungal pathogens, and as a result >50% of infected patients do not survive [7, 8]. Glucocorticoids can inhibit multiple different molecular and cellular pathways, and it is not clear which of these effects is the main cause of susceptibility to invasive aspergillosis and other opportunistic infections. This knowledge is necessary to develop novel therapeutic approaches to treat patients with invasive aspergillosis who are undergoing glucocorticoid therapy or to develop safe glucocorticoid therapy with a lower risk of opportunistic infection.

Glucocorticoids exert their activity by binding to the glucocorticoid receptor (GR) which is a nuclear receptor [2]. Upon binding to ligand in the cytosol, GR translocates to the nucleus and activates or represses gene transcription [2]. GR can affect gene expression through three mechanisms: directly binding to glucocorticoid response elements (GRE) in the DNA sequence, trans-repression through binding to other transcription factors, or composite binding to both GREs and transcription factors at the same time [2, 3]. It is thought that the immunosuppressive effects of glucocorticoids are mainly mediated by trans-repression of nuclear factor-κB (NF-κB) [1, 9–11]. NF-κB is a family of transcription factors, including the canonical p65 and p50 subunits, that activate inflammatory responses by promoting transcription of various signaling molecules such as cytokines [12]. Under resting conditions, p65/p50 heterodimers are bound by an inhibitor, IκB, and sequestered in the cytoplasm [13–17]. Downstream of activation of pattern recognition receptors (PRRs), cytokine receptors, or T/B-cell receptors [18, 19], IκB is phosphorylated by the multi-subunit IκB kinase (IKK) complex, which is composed of IKKα, IKKβ, and a regulatory subunit IKKγ (NEMO) [17–20]. This phosphorylation leads to degradation of IκB, releasing the NF-κB dimers which rapidly translocate to the nucleus to initiate target gene expression [12, 17–19]. To inhibit this activation, GR can both directly bind and trans-repress NF-κB subunits and directly bind to the IκB gene promoter to induce transcription. The relative significance of each of these activities in NF-κB suppression by GR is debated [21–23]. Additionally, if glucocorticoid-mediated suppression of NF-κB is the major mechanistic cause of susceptibility to invasive aspergillosis, rather than GR-mediated effects on other pathways, is not well understood. *A. fumigatus* can induce NF-κB activation *in vitro* in monocytes, in bronchial epithelial cells in mice, and at the site of infection in larval zebrafish [24–26]. This activation is likely required for activation of the innate immune system, including macrophages and neutrophils, the first line of defense against *A. fumigatus* infection. However, how NF-κB activation regulates macrophage- and neutrophil-mediated control of *A. fumigatus* and if glucocorticoids suppress these control mechanisms *in vivo* is not understood.

In this study, we investigate these questions in a zebrafish larva-*Aspergillus* infection model which allows us to non-invasively image this dynamic host-pathogen interaction inside of a live, intact host over multiple days. Using this model, we have previously shown that macrophages mainly respond to *A. fumigatus* spores and prevent germination, while neutrophils are only recruited after spore germination into hyphae [27–29]. This is in line with previous findings that macrophages efficiently kill spores and neutrophils are efficient at killing hyphae in cell culture [30–32]. Consistently, in humans and in mammalian models, alveolar macrophages are likely to encounter spores and neutrophils are recruited from the blood secondarily [33]. The immune system of zebrafish is largely conserved with humans and during the larval stage primarily consists of macrophages and neutrophils, facilitating the study of phagocyte-specific mechanisms with no interference from the adaptive system [34]. The zebrafish model has been instrumental in modeling a range of human infections to better understand host-pathogen mechanisms, such as mycobacterial granuloma formation [35].

We report that exposure to the glucocorticoid drug dexamethasone significantly increases the mortality of *A. fumigatus*-infected larvae, recapitulating the susceptibility of glucocorticoid-treated human patients. Through CRISPR/Cas9 targeting of GR, we demonstrate that dexamethasone activity is mediated via GR in larval zebrafish. To determine the host innate mechanisms that are inhibited by dexamethasone treatment we used daily, live imaging of infected larvae in combination with established innate immune cell-deficiency models. We report that the increased mortality of infected hosts is primarily due to a decrease in neutrophil-mediated control of invasive hyphal growth. Dexamethasone treatment induces IκB transcription and suppresses *A. fumigatus*-induced NF-κB activation. Although inhibition of other minor pathways may also promote susceptibility to *A. fumigatus* infection, CRISPR/Cas9 targeting of IKKγ phenocopies dexamethasone treatment, suggesting that the effects of glucocorticoids are largely due to the inhibition of NF-κB signaling and inhibition of neutrophil function.

## Results

### Dexamethasone exposure decreases survival of zebrafish larvae infected with *A. fumigatus* and suppresses NF-κB activation at the infection site

Glucocorticoid drugs such as dexamethasone increase host susceptibility to *A. fumigatus* infection, but the cellular mechanisms through which this occurs are largely unknown [6]. To investigate this question, we used an established larval zebrafish host model [26, 27, 36]. We injected *A. fumigatus* spores of the CEA10 strain into the hindbrain ventricle of 2 days post fertilization (dpf) wild-type larvae and immediately exposed larvae to 10 μM dexamethasone or a DMSO vehicle control. Dexamethasone-exposed larvae succumb to the infection at a significantly higher rate than control larvae, with a hazard ratio of 2.5, indicating that dexamethasone-exposed larvae are 2.5 times more likely to succumb to the infection as compared to control larvae (Fig 1A), consistent with previous results [26]. No significant survival defect due to dexamethasone treatment is observed in PBS-injected mock-infected larvae (Fig 1A).

**Figure 1.**
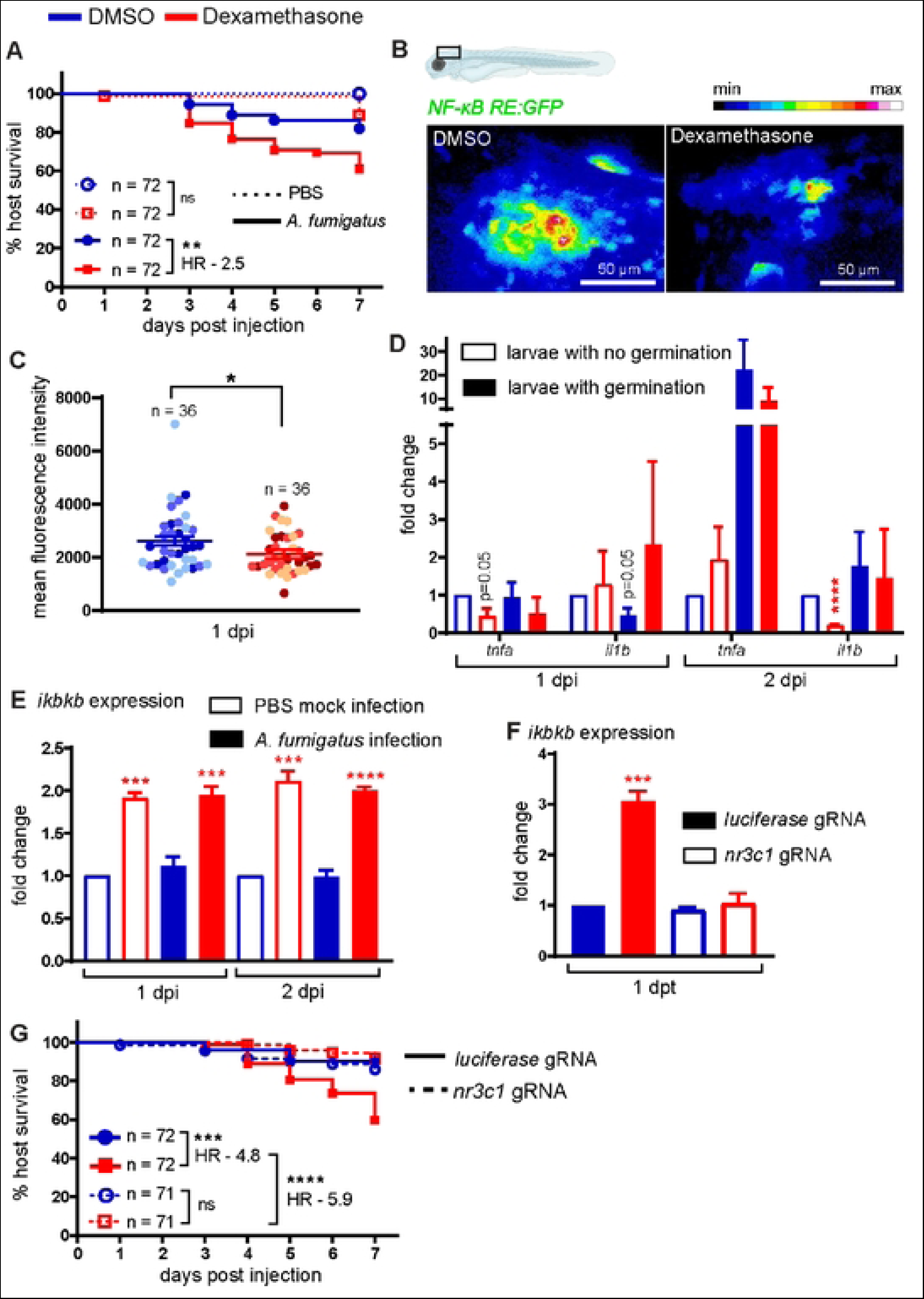
Dexamethasone suppresses NF-κB activation and increases susceptibility to *Aspergillus fumigatus* via the glucocorticoid receptor. **(A)** Survival of wild-type larvae injected at 2 dpf with CEA10 *A. fumigatus* spores or PBS mock-infection in the presence of 10 μM dexamethasone or DMSO vehicle control. At least 24 larvae per condition, per replicate were used and the total larval N per condition is indicated. Cox proportional hazard regression analysis was used to calculate P values and hazard ratio (HR). Average injection CFUs: dexamethasone = 15, DMSO = 12. **(B, C)** Larvae of NF-κB reporter line (*Tg(NF-κB RE:GFP))* were injected with CEA10 *A. fumigatus* spores and were exposed to 10 μM dexamethasone or DMSO. Larvae were imaged at 1 dpi. (B) Representative images showing relative GFP expression from z projection of 6 slices. Scale bar = 50 μm. (C) Quantification of fluorescent expression in the hindbrain ventricle at 1 dpi is shown with emmeans ± SEM from three independent replicates and the total larval N per condition is indicated. Each data point represents an individual larva, color-coded by replicate. P values were calculated by ANOVA. **(D)** Larvae were injected with GFP-expressing TFYL49.1 (CEA10) spores and exposed to 10 μM dexamethasone or DMSO. At 1 and 2 dpi, larvae were screened for germination and total RNA was extracted from each pooled group. RT-qPCR analysis of cytokine expression in pooled larvae is shown. Data are normalized to DMSO no germination control group. P values were calculated by Student’s t-test. Data are from three independent replicates. **(E)** NF-κB inhibitor *ikbkb* expression in larvae injected with CEA10 spores or PBS mock-infection and exposed to 10 μM dexamethasone or DMSO is shown. Total RNA was extracted at 1 and 2 dpi from 20 pooled larvae per condition per day. Data are normalized to DMSO PBS mock-infection at each day post injection. P values were calculated by Student’s t-test. Data are from three independent replicates. **(F)** Embryos at 1 cell stage were injected with gRNAs targeting glucocorticoid receptor gene *nr3c1* or *luciferase* control together with Cas9 protein. At 2 dpf, larvae were treated with 10 μM dexamethasone or DMSO. Total RNA from 20 pooled larvae per condition was extracted at 1 day post treatment (dpt) and *ikbkb* expression was quantified using RT-qPCR. Data are normalized to *luciferase* gRNA + DMSO group. P values were calculated by Student’s t-test. Data are from three independent replicates. **(G)** Survival of *nr3c1* mutant larvae injected with CEA10 spores and exposed to 10 μM dexamethasone or DMSO. Data are pooled from three independent replicates, at least 23 larvae per condition, per replicate and the total larval N per condition is indicated. Cox proportional hazard regression analysis was used to calculate P values and hazard ratios (HR). Average injection CFUs: *nr3c1* = 25 or *luciferase* = 26.

The *A. fumigatus* CEA10 strain induces NF-κB activation at the site of infection, and glucocorticoids can suppress NF-κB activity [1, 26]. To test if dexamethasone suppresses NF-κB activation in this infection model, we used a previously published NF-κB reporter transgenic zebrafish line that express EGFP under an NF-κB responsive promoter (*Tg(NF-κB RE:GFP)*) [37]. We injected *NF-κB RE:GFP* larvae with *A. fumigatus* CEA10 spores, imaged the infection site 1 and 2 days post injection (dpi), and quantified EGFP expression (Fig 1B). Dexamethasone-exposed larvae have lower EGFP expression than control larvae, although this difference is only statistically significant at 1 dpi (Figs 1C and S1A). We then tested if dexamethasone suppresses the expression of specific NF-κB target cytokine genes using RT-qPCR. Since fungal germination drives NF-κB activation we screened larvae by microscopy prior to RNA extraction and split larvae into two groups based on whether hyphae were present or not. At 1 dpi, the expression of NF-κB target genes *tnfa* and *il1b* was not yet induced by germination, although dexamethasone treatment significantly inhibited *tnfa* expression even in larvae without germination (Fig 1D). At 2 dpi, germination increases *tnfa* expression ~20-fold in larvae exposed to DMSO, and this is only partially suppressed by dexamethasone treatment, potentially because dexamethasone treated larvae may experience more hyphal growth and therefore more immune activation overall (Fig 1D). *il1b* is only induced ~2-fold by germination at 2 dpi but dexamethasone significantly inhibits *il1b* expression in larvae without germinated spores (Fig 1D). Another marker of macrophage activation, *irg1*, is also induced by germination at 1 dpi and inhibited by dexamethasone treatment (S1B Fig). Additionally, the expression of anti-inflammatory genes *il10* and *tgfb* are significantly inhibited by dexamethasone treatment at 1 dpi but increased at 2 dpi (S1B Fig). Overall, these data demonstrate that dexamethasone can affect host gene expression of inflammatory markers, including NF-κB-regulated genes, although increased fungal germination and growth may override this suppression.

To determine the mechanism through which dexamethasone inhibits NF-κB activation, we tested the expression of the *ikbaa* gene which encodes IκBα, the inhibitor of NF-κB. Dexamethasone-treated larvae have higher expression of *ikbaa*, regardless of infection status (Fig 1E). These results demonstrate that during *A. fumigatus* infection, one mechanism through which glucocorticoids inhibit NF-κB activation is by increasing transcription of this inhibitor.

### Glucocorticoid receptor is required for dexamethasone-mediated immunosuppression

In mammals, glucocorticoids primarily mediate their effects through the glucocorticoid receptor [38]. To determine if the increased susceptibility of dexamethasone-treated larvae to *A. fumigatus* infection was due to signaling through this receptor, and not to off-target effects on either the host or pathogen, we used CRISPR/Cas9 to target the zebrafish glucocorticoid receptor gene *nr3c1*. We designed two gRNAs: one targeting exon 2 which encodes the N-terminal domain and the other targeting exon 4 which encodes part of the DNA binding domain (S2A Fig). We injected embryos at the 1 cell stage with both gRNAs targeting *nr3c1* or control gRNAs targeting *luciferase*, together with Cas9 protein. PCR using primers flanking the target sites on genomic DNA isolated from 2 dpf larvae confirmed successful targeting of DNA (S2B Fig). In these same F0 injected crispants, we tested if dexamethasone can induce *ikbaa* expression as seen with wild-type larvae (Fig 1E). Dexamethasone significantly induces *ikbaa* expression in control larvae but fails to induce any expression in *nr3c1* crispant larvae (Fig 1F), indicating that GR function is abolished in these crispant larvae and that IκB-mediated suppression of NF-κB activation by dexamethasone depends on the glucocorticoid receptor. Further, we tested the effects of *nr3c1* mutation in survival of infected larvae. While dexamethasone-exposed control larvae succumb to *A. fumigatus* infection, dexamethasone has no effect on survival of infected *nr3c1* crispant larvae (Fig 1G). Targeting of *nr3c1* had no effect on the survival of PBS mock-infected larvae (S2C Fig). Additionally, direct exposure of *A. fumigatus* spores to dexamethasone has no effect on spore germination or hyphal growth (S3 Fig). These data indicate that the immunosuppressive effects of dexamethasone in the context of *A. fumigatus* infection depend solely on signaling through a functional glucocorticoid receptor.

### Dexamethasone partially suppresses macrophage recruitment, but not neutrophil recruitment

We next sought to understand how dexamethasone mediates phagocyte responses to *A. fumigatus*. As dexamethasone suppresses pro-inflammatory cytokine expression (Fig 1D), we hypothesized that phagocyte recruitment would be inhibited by dexamethasone. We injected ~30 GFP-expressing *A. fumigatus* spores into larvae expressing mCherry in macrophages (*Tg(mpeg1:H2B-mCherry)*) and BFP in neutrophils (*Tg(lyz:BFP)*) and treated larvae with dexamethasone or DMSO vehicle control. We enumerated macrophage and neutrophil recruitment to the infection site through daily, live confocal imaging of infected larvae. In line with previous findings [26, 39], macrophages arrive first and form clusters around spores starting at 1 dpi (Fig 2A). A significantly higher number of macrophages arrive at 2 dpi in control larvae compared to dexamethasone-treated larvae, yet ~90 macrophages still arrive at the infection site in dexamethasone-treated larvae (Fig 2B). Macrophage cluster area is not significantly different between the two groups (Fig 2C). Macrophage clusters resolve from 3-5 dpi in DMSO-exposed larvae (Fig 2B, C). However, in dexamethasone-exposed larvae, more macrophages are recruited later in the infection with a significantly higher number of macrophages and larger cluster area at 5 dpi (Fig 2B, C). Neutrophils respond starting at 2 dpi, primarily after spores start to germinate, and neutrophils are able to infiltrate into macrophage clusters (Fig 2A). The number of recruited neutrophils is not significantly different between dexamethasone- and DMSO-treated larvae at 1, 2, or 3 dpi (Fig 2D). At 5 dpi, similar to macrophages, a significantly higher number of neutrophils is present at the infection site in larvae exposed to dexamethasone compared to control larvae (Fig 2D). This is likely due to increased fungal growth attracting more macrophages and neutrophils to the infection site, as described previously [39]. Injected *A. fumigatus* spores can germinate, and after germination, the fungal filamentous growth can branch and form a network of hyphae, which we classified as invasive hyphae. To normalize for differences in this fungal growth, we analyzed the number of macrophages and neutrophils at specific stages of fungal germination and invasive hyphal growth. On the day that germination is first observed in larvae, the number of macrophages at the infection site is significantly lower in dexamethasone-exposed larvae compared to control larvae (Fig 2E). However, macrophage numbers on the day before germination is observed and on the day that invasive hyphae is first observed are not significantly different between dexamethasone- and control-treated larvae (Fig 2E), and neutrophil numbers are also similar between the two conditions throughout the infection (Fig 2F). Overall, we find that many phagocytes are still recruited to the infection site in dexamethasone-treated larvae. Dexamethasone has no effect on neutrophil recruitment but can curb macrophage recruitment earlier in infection. However, immune activation from fungal germination can override dexamethasone-mediated suppression of macrophage recruitment at later stages of infection.

**Figure 2.**
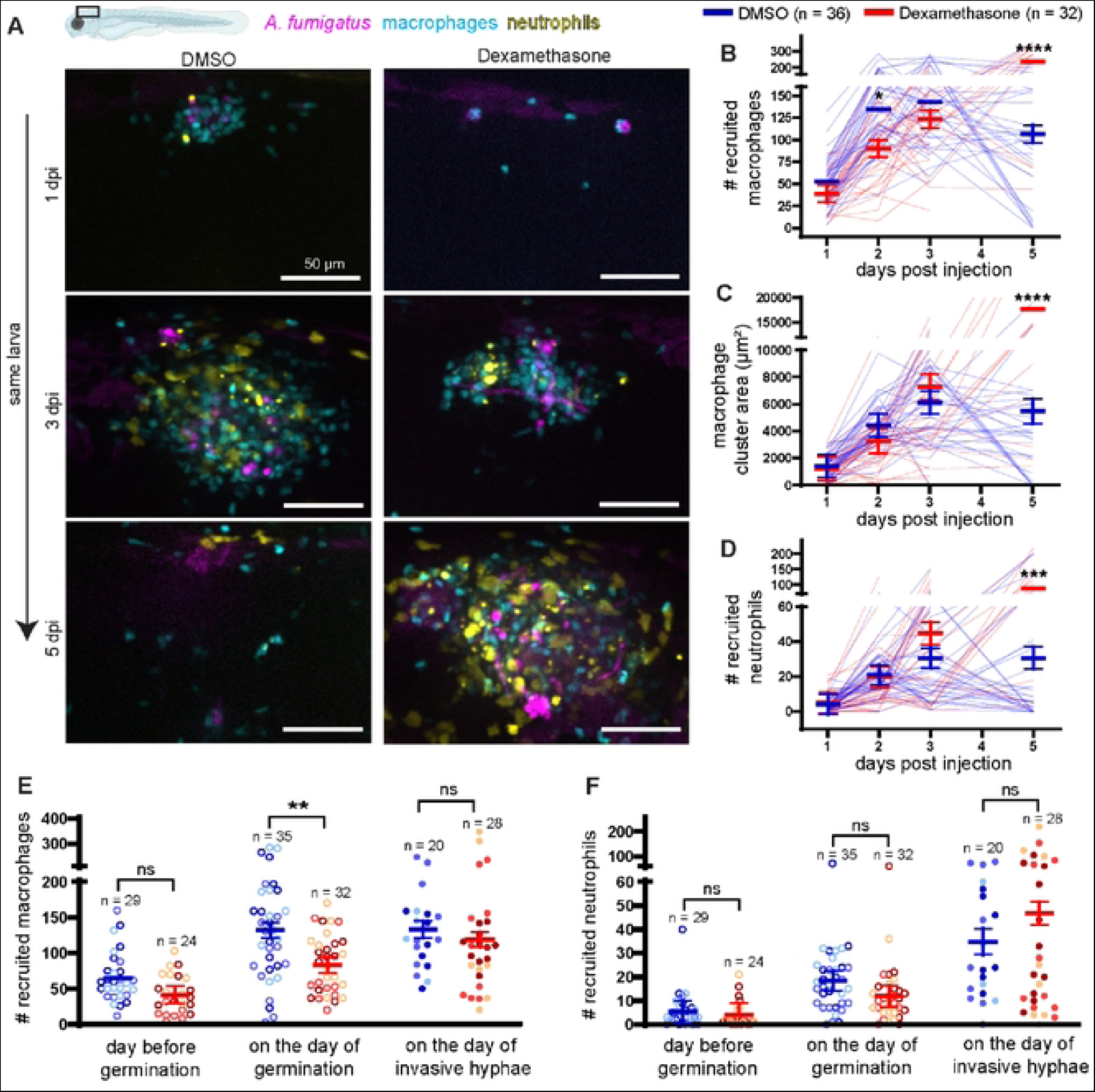
Dexamethasone moderately suppresses macrophage recruitment but not neutrophil recruitment. Larvae with labeled macrophages (*Tg(mpeg1:H2B-mCherry)*) and neutrophils (*Tg(lyz:BFP)*) were injected with GFP-expressing TFYL49.1 (CEA10) spores at 2 dpf, exposed to 10 μM dexamethasone or DMSO vehicle control and live imaged at 1, 2, 3, and 5 dpi. Data are pooled from three independent replicates, at least 10 larvae per condition, per replicate. **(A)** Representative images show different patterns of phagocyte recruitment across multiple days in larvae exposed to dexamethasone or DMSO. Scale bar = 50 μm. **(B)** Number of macrophages recruited, **(C)** macrophage cluster area, and **(D)** number of neutrophils recruited were quantified from the images. (B-D) Bars represent emmeans ± SEM and P values were calculated by ANOVA. Each line represents an individual larva. **(E, F)** Number of recruited macrophages (E) and neutrophils (F) one day before germination occurred, on the day of germination, and on the day invasive hyphae occurred were plotted for larvae that experienced fungal growth. Bars represent emmeans ± SEM and P values were calculated by ANOVA. Each data point represents an individual larva, color-coded by replicate.

### Spore killing is not significantly impacted by dexamethasone exposure

As phagocytes are able to migrate to the site of infection even in dexamethasone-treated larvae, we hypothesized that the functions of phagocyte-mediated control of *A. fumigatus* are impacted by dexamethasone. The first step in the phagocyte response is macrophage-mediated phagocytosis of spores and spore killing [33, 40]. To test whether dexamethasone treatment decreases macrophage-mediated spore killing, we used an established live-dead staining method in which *A. fumigatus* spores expressing GFP are coated with AlexaFluor546 via cell wall cross-linking [26, 39]. We injected these spores into larvae expressing BFP in macrophagesm (*Tg(mfap4:BFP)*) and imaged the infection site at 2 dpi. We then quantified the percentage of live spores (AlexaFluor+ and GFP+) versus dead spores (AlexaFluor+ and GFP-) (Fig 3A). Dexamethasone-treated larvae are slightly worse at killing injected spores than control larvae, although this difference is not statistically significant (Fig 3B). To further quantify spore burden over time, we homogenized and plated larvae to quantify CFUs from dexamethasone-treated or control larvae across 7 days of infection. In agreement with the live-dead staining results, we find no significant difference in CFU burden in dexamethasone-treated larvae compared to control larvae (Fig 3C).

**Figure 3.**
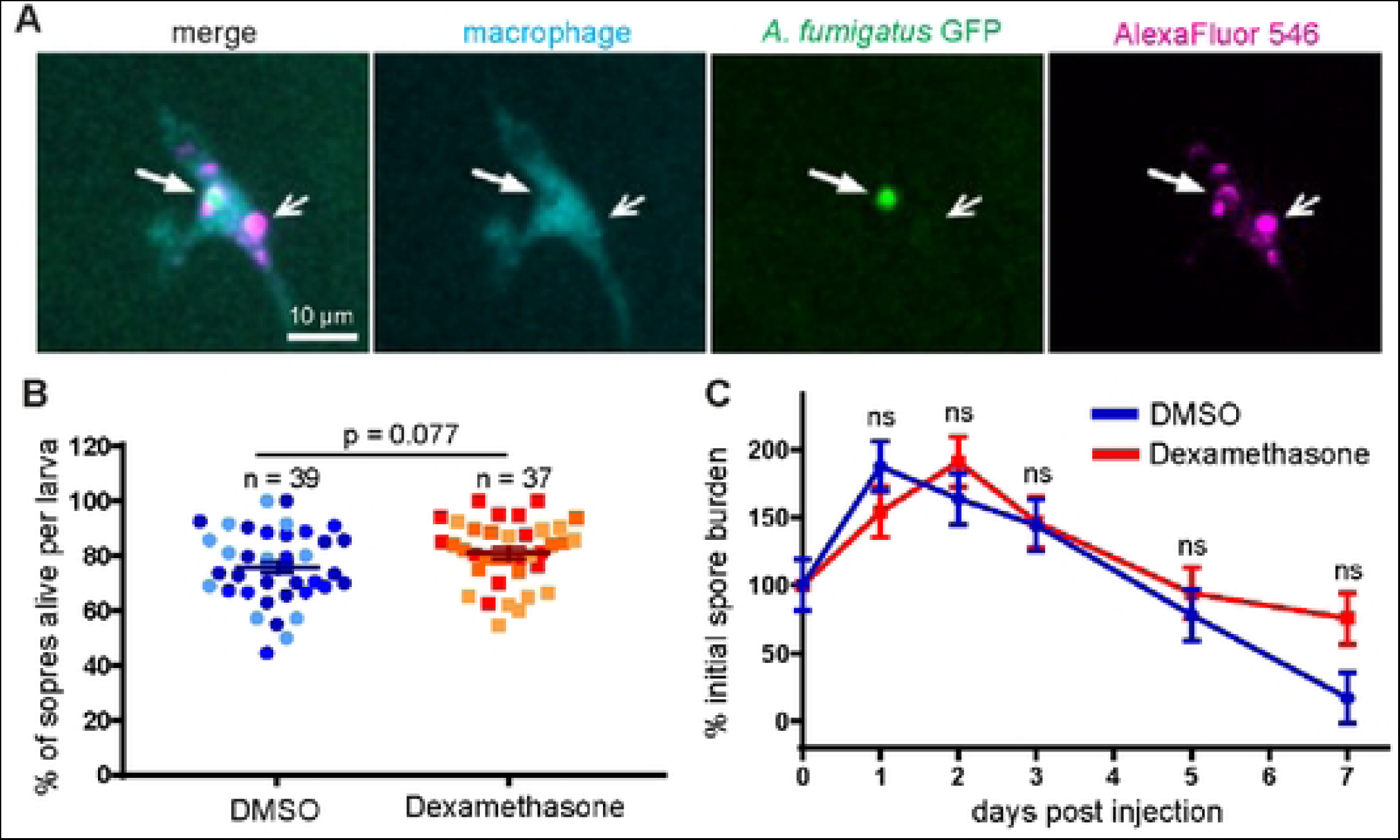
Dexamethasone does not affect spore killing. **(A, B)** Macrophage-labeled larvae (*Tg(mfap4:BFP)*) were injected with GFP-expressing *A. fumigatus* TFYL49.1 (CEA10) spores coated with AlexaFluor546 at 2 dpf, exposed to 10 μM dexamethasone or DMSO vehicle control, and live imaged at 2 dpi. (A) Representative images of z projection of 3 slices showing live (filled arrow) and dead (open arrow) spores within a macrophage. Scale bar = 10 μm. (B) The percentage of live spores in the hindbrain per larvae is shown with bars representing emmeans ± SEM from three independent replicates, and the total larval N per condition is indicated. Each data point represents an individual larva, color-coded by replicate. P values were calculated by ANOVA. **(C)** Wild-type larvae were injected with CEA10 spores at 2 dpf, exposed to 10 μM dexamethasone or DMSO vehicle control, and fungal burden was quantified by homogenizing and plating individual larvae for CFUs at multiple days post injection. Eight larvae per condition, per dpi, per replicate were quantified, and the number of CFUs at each dpi is represented as a percentage of the initial spore burden. Bars represent emmeans ± SEM from three independent replicates, P values calculated by ANOVA. Average injection CFU: 32.

### Invasive hyphal growth post-germination is increased in dexamethasone-treated larvae

As spore killing is only minorly inhibited by dexamethasone treatment, we hypothesized that in dexamethasone-exposed hosts phagocytes fail to control *A. fumigatus* germination and invasive hyphal growth. To quantify fungal growth over time, we went back to our daily imaging experiment (Fig 2) and monitored spore germination and invasive hyphal growth from the GFP signal expressed by *A. fumigatus*. As these experiments were done with a fast-germinating CEA10-derived strain, spore germination occurs at high levels by 2 dpi and the rate of spore germination is not significantly different in dexamethasone-exposed larvae compared to control DMSO-exposed larvae (Fig 4A, B). The cumulative percentage of larvae that experience invasive hyphal growth (presence of branched hyphae) is significantly higher with dexamethasone exposure compared to control conditions (Fig 4B). We also quantified the fungal burden in larvae across the full 5 day experiment by measuring the GFP+ area, confirming that dexamethasone-treated larvae experience significantly more fungal growth compared to control larvae (Fig 4C). Next, we rated the severity of fungal growth using a scoring system of 0 to 4, from no germination to invasive hyphal growth (S4 Fig). In larvae exposed to dexamethasone, invasive hyphal growth becomes severe quickly, within 2-3 days, and causes mortality, while many control larvae are able to delay this invasive growth (Fig 4D). These data suggest that dexamethasone treatment decreases the ability of host immune cells to inhibit post-germination invasive hyphal growth of *A. fumigatus*.

**Figure 4.**
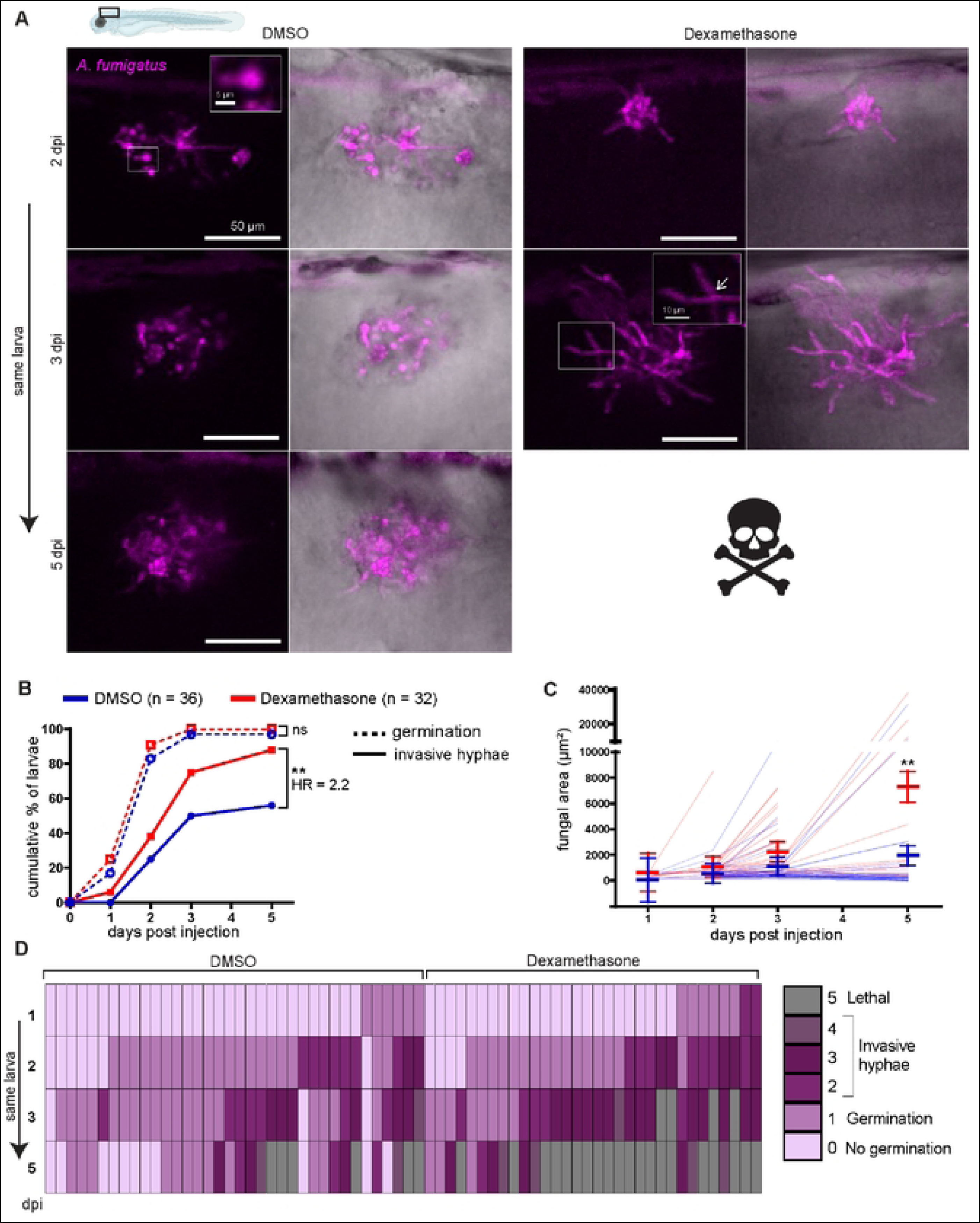
Dexamethasone suppresses immune control of *A. fumigatus* invasive hyphal growth. Wild-type larvae were injected with GFP-expressing TFYL49.1 (CEA10) spores at 2 dpf, exposed to 10 μM dexamethasone or DMSO vehicle control and imaged at 1, 2, 3, and 5 dpi. Data are pooled from three independent replicates, at least 10 larvae per condition, per replicate. **(A)** Representative images show hyphal growth differences in larvae exposed to dexamethasone or DMSO. Insets show a germinated spore and branched invasive hyphae (open arrow). Scale bar = 50 μm (5 and 10 μm in insets). **(B)** Cumulative percentage of larvae with germination (dotted line) and invasive hyphae (solid line) through 5 dpi. Cox proportional hazard regression analysis was used to calculate P values and hazard ratios (HR). **(C)** In larvae with fungal germination, fungal area was quantified from maximum intensity projection images. Each line represents an individual larva and bars represent emmeans ± SEM. **(D)** Severity of fungal growth was scored for all larvae and displayed as a heatmap. Representative images for each score can be found in S4 Fig.

### Dexamethasone predominantly suppresses neutrophil function to cause host susceptibility

To confirm that dexamethasone impacts the ability of host innate immune cells to control the invasive growth stages of *A. fumigatus*, we generated zebrafish larvae without macrophages and neutrophils, the primary innate immune cells active in zebrafish larvae[34]. If dexamethasone increases host susceptibility by inhibiting the function of these cells, then in larvae that already lack these cells we expect that dexamethasone treatment would not significantly decrease host survival. To prevent the development of both macrophages and neutrophils we injected 1 cell embryos with a high concentration of *pu.1* morpholino, knocking down expression of the *pu.1* (*spi1b*) transcription factor required for phagocyte development [41]. In larvae that do not develop phagocytes, >90% of larvae succumb to *A. fumigatus* infection regardless of dexamethasone exposure, supporting the idea that dexamethasone primarily inhibits the function of these cells to cause host susceptibility (S5A Fig).

While neutrophil function is thought to predominate during control of invasive growth post-germination, macrophages can also attack hyphae and may play a role in control of hyphal growth [42]. We therefore sought to determine the relative impact of dexamethasone on macrophage versus neutrophil function against infection. To do this, we employed established models of neutrophil-defective or macrophage-deficient larvae. If dexamethasone predominantly suppresses neutrophil-mediated mechanisms, we expect dexamethasone to cause no additional survival defect in larvae that already do not have functional neutrophils. We infected neutrophil-defective (*Tg(mpx:mCherry-2A-rac2D57N)*) larvae, in which neutrophils are unable to migrate to the infection site, with *A. fumigatus* spores, treated larvae with either dexamethasone or DMSO vehicle control, and monitored survival [43]. Consistent with previous results [26], neutrophil-defective larvae succumb to the infection at a higher rate than control wild-type larvae (Fig 5A). Dexamethasone further decreases survival of neutrophil-defective larvae (Fig 5A). While wild-type larvae treated with dexamethasone are 4.3 times more likely succumb to the infection compared to vehicle control larvae, neutrophil-defective larvae are only 2.2 times more likely to succumb due to treatment. Dexamethasone treatment does not affect survival of PBS mock-infected larvae (S5B Fig). These results suggest that dexamethasone can affect the function of other cell types remaining in these neutrophil-defective larvae but that neutrophils are also a major target of dexamethasone.

**Figure 5.**
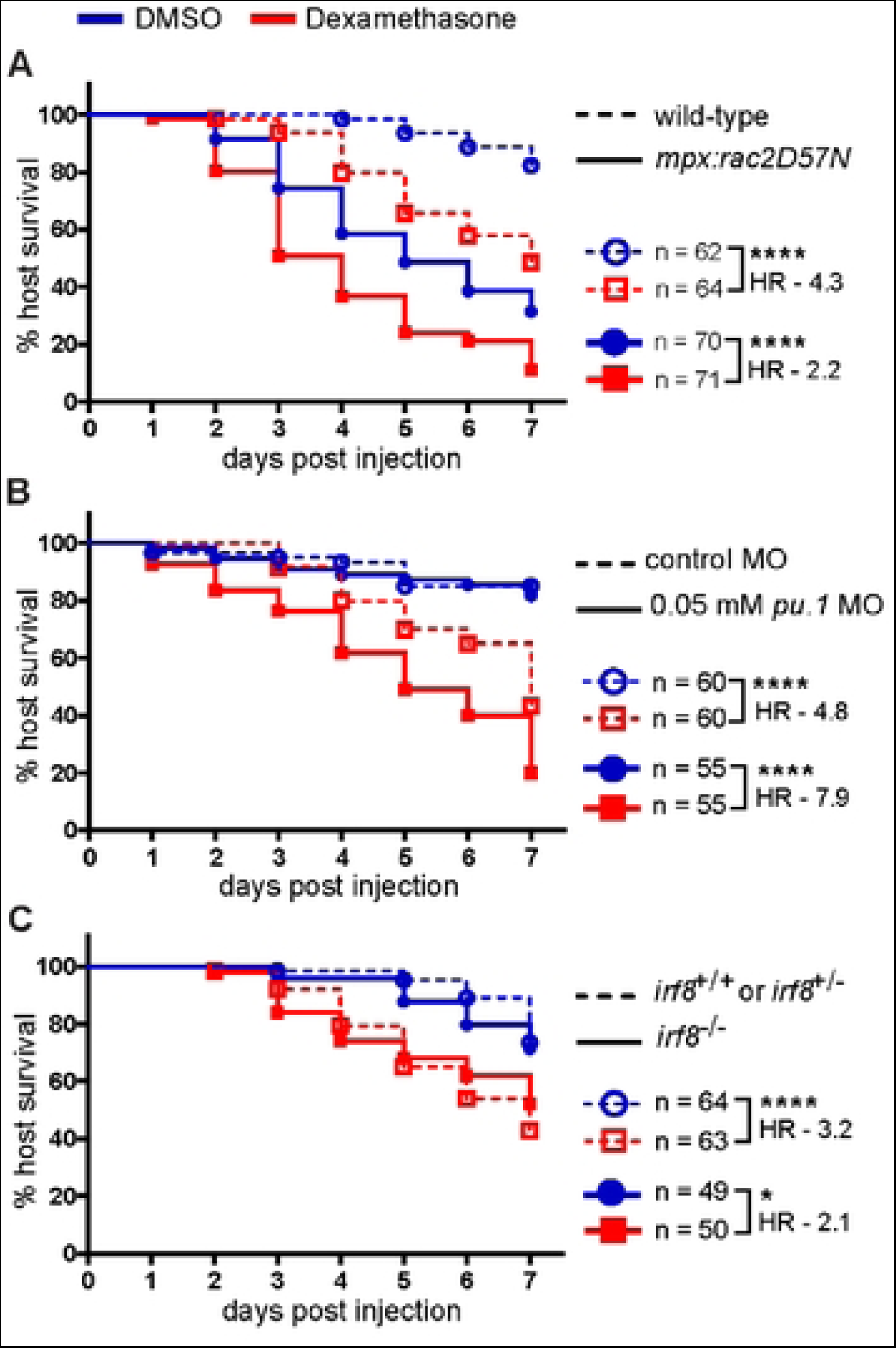
Dexamethasone primarily suppresses neutrophil-mediated host protection. Survival of larvae injected at 2 dpf with CEA10 *A. fumigatus* spores and exposed to 10 μM dexamethasone or DMSO vehicle control. Data are pooled from three independent replicates, at least 12 larvae per condition, per replicate and the total larval N per condition is indicated in each figure. Cox proportional hazard regression analysis was used to calculate P values and hazard ratios (HR). **(A)** Survival of neutrophil-defective larvae (*mpx*:*rac2D57N*) and wild-type larvae. Average injection CFUs: wild-type = 26, *mpx*:*rac2D57N* = 29. **(B)** Survival of macrophage-deficient or control larvae. Development of macrophages was inhibited by 0.05 mM *pu.1* morpholino (MO). Control larvae received standard control MO. Average injection CFUs: control MO = 31, *pu.1* MO = 25. **(C)** Survival of macrophage-deficient *irf8^−/−^* or control (*irf8^+/+^* or *irf8^+/−^*) larvae. Average injection CFUs: *irf8^+/+^* or *irf8^+/−^*= 71, *irf8^−/−^* = 50.

Next, we performed these experiments in macrophage-deficient larvae. First, we used clodronate liposomes to deplete macrophages. Macrophage-depleted larvae succumb to the infection, and dexamethasone further increases mortality of these larvae (S5C Fig). However, dexamethasone causes significant mortality in macrophage-depleted PBS-injected mock-infected larvae (S5D Fig), making results from these experiments difficult to interpret. Instead, to prevent the development of macrophages, we used a low concentration of *pu.1* morpholino that affects macrophage but not neutrophil development [44]. We confirmed that at this concentration macrophages are depleted but neutrophils remain intact and functional (S5E-G Fig). Consistent with previous results [26], macrophage deficiency alone does not significantly affect the survival of larvae infected with the CEA10 strain of *A. fumigatus* (Fig 5B). Additionally, dexamethasone does not affect the survival of PBS mock-infected *pu.1* morphant larvae (S5H Fig). However, dexamethasone exposure increases the susceptibility of both control and macrophage-deficient *A. fumigatus* infected larvae (Fig 5B). Dexamethasone-exposed macrophage-deficient larvae are 7.9 times more likely to succumb to the infection compared to vehicle exposure, while control larvae which are 4.8 times more likely to succumb to the infection after dexamethasone treatment (Fig 5B). Therefore, when macrophages are not present and neutrophils are the major immune cell present, dexamethasone has a greater impact on host survival. These results are in contrast to experiments in the neutrophil-defective line, where macrophages are the major immune cell present and dexamethasone has a lower impact on host survival. Overall, these data demonstrate that dexamethasone-mediated suppression of neutrophil function is a major cause of host susceptibility to *A. fumigatus* infection.

To further study the effect of dexamethasone on neutrophil function, we used an *irf8* mutant line, which lacks macrophages and has a compensating increase in neutrophil numbers [45]. *irf8^−/−^* larvae are significantly more susceptible to infection when exposed to dexamethasone compared to vehicle control (Fig 5C). However, the relative increase in susceptibility is similar to that of *irf8^+/−^* and *irf8^+/+^* control larvae, suggesting that an increased number of neutrophils can partially compensate for the suppressive effects of dexamethasone on neutrophil function (Fig 5C).

### Neutrophils fail to control invasive hyphal growth in dexamethasone-treated larvae

So far, we have established that dexamethasone predominantly suppresses the function of neutrophils to cause susceptibility to *A. fumigatus* (Fig 5). This is consistent with our finding that dexamethasone exposure leads to increased invasive hyphal growth inside of larvae (Fig 4), as neutrophils are the primary innate immune cell responsible for clearing *A. fumigatus* hyphae. To focus specifically on the effects of dexamethasone on neutrophil-mediated control of hyphae, we decided to further characterize the effects of dexamethasone exposure on infected *irf8^−/−^* larvae which lack macrophages and have an excess of neutrophils. We infected *irf8^−/−^* larvae with labeled neutrophils (*Tg(lyz:BFP)*) with a GFP-expressing strain of *A. fumigatus*, treated larvae with dexamethasone or vehicle control, and performed live imaging at 1, 2, 3, and 5 dpi (Fig 6). As none of these larvae have macrophages which are the cell type primarily responsible for preventing spore germination [26], we observed similar levels of germination in larvae exposed to both dexamethasone and DMSO, with all larvae harboring germinated spores by 3 dpi (Fig 6A, B). However, the cumulative percentage of larvae harboring invasive hyphae is significantly higher in dexamethasone-treated larvae compared to vehicle control-treated larvae (Fig 6B). This is due to a lack of fungal growth control post-germination. There is no significant effect of dexamethasone treatment on the day that germination occurs, with the majority of larvae experiencing germination at 1 dpi regardless of treatment group (Fig 6D). However, dexamethasone-exposed larvae develop initial invasive hyphae as early as ~2 dpi on average, while it takes ~3 days for DMSO-exposed larvae to develop invasive hyphae (Fig 6E). Once germination occurs, invasive hyphae appears on average 1 day later in dexamethasone-exposed larvae as compared to an average of 1.5 days later in DMSO-exposed larvae (Fig 6F). Analysis of the severity of fungal growth or quantification of fungal area in all individual larvae across all days of the experiment demonstrates that dexamethasone-treated *irf8* mutant larvae experience uncontrolled hyphal growth (Figs 6C and S6A). Although germination occurs in all larvae, neutrophils in DMSO-treated larvae are able to delay the development of severe invasive growth of hyphae, while dexamethasone-exposed larvae quickly develop severe invasive hyphae and succumb to the infection (Fig 6C).

**Figure 6.**
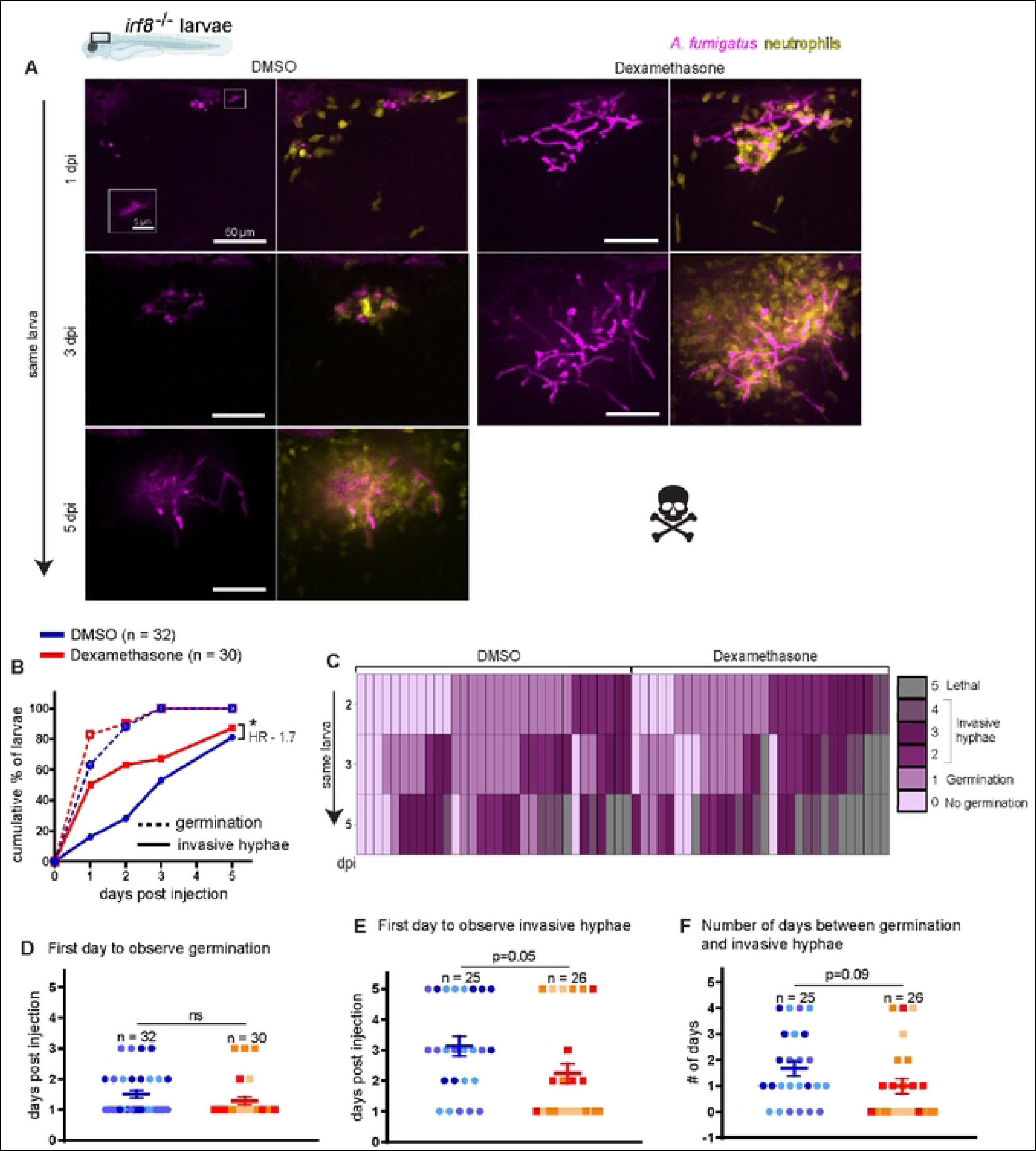
Dexamethasone suppresses neutrophil-mediated control of *A. fumigatus* invasive hyphal growth. Macrophage-deficient *irf8^−/−^* larvae with labeled neutrophils (*Tg(lyz:BFP)*) were injected with GFP-expressing TFYL49.1 (CEA10) spores at 2 dpf, exposed to 10 μM dexamethasone or DMSO vehicle control and live imaged at 1, 2, 3, and 5 dpi. Data are pooled from three independent replicates, at least 10 larvae per condition, per replicate. **(A)** Representative images show hyphal growth differences and neutrophil recruitment in larvae exposed to dexamethasone or DMSO. Inset shows a germinated spore. Scale bar = 50 μm (inset 5 μm). **(B)** Cumulative percentage of larvae with germination (dotted line) and invasive hyphae (solid line) through 5 dpi. Cox proportional hazard regression analysis was used to calculate P values and hazard ratios (HR). **(C)** Severity of fungal growth was scored for all larvae and displayed as a heatmap. Representative images for each score can be found in S4 Fig. **(D-F)** In larvae in which fungal growth occurred, the day on which germination (D) and invasive hyphae (E) was first observed and the number of days between germination and invasive hyphae (F) are plotted. Bars represent emmeans ± SEM and P values were calculated by ANOVA. Each data point represents an individual larva, color-coded by replicate.

In dexamethasone-treated *irf8^−/−^* larvae, we observed many neutrophils at the infection site, clustering around the fungus (Fig 6A), suggesting that in these larvae dexamethasone inhibits neutrophil function rather than neutrophil recruitment, as we observed in wild-type larvae (Fig 2). To confirm this, we quantified neutrophil cluster area both across all days of imaging and normalized to the germination status of larvae, finding no statistically significant differences between larvae treated with dexamethasone or vehicle control (S6B, C Fig). As expected, there is a positive correlation between fungal area and neutrophil cluster area in both vehicle- and dexamethasone-treated larvae (S6D Fig). However, when we plotted the neutrophil cluster area relative to the fungal area, the slope of this correlation is slightly lower in dexamethasone-treated larvae, suggesting that for a given fungal load these larvae may recruit fewer neutrophils (S6D Fig). Overall, a decrease in neutrophil recruitment may play a minor role in the lack of fungal control in *irf8* mutant dexamethasone-treated larvae, but an inhibition of neutrophil function against hyphae by this drug is likely the major factor leading to host susceptibility.

### Neutrophil-mediated control of invasive hyphal growth requires NF-κB

Our results thus far demonstrate that 1) dexamethasone treatment inhibits NF-κB at the site of *A. fumigatus* infection and 2) dexamethasone primarily impacts neutrophil function to cause host susceptibility to *A. fumigatus*. We therefore wanted to confirm that NF-κB signaling is required for neutrophil function against *A. fumigatus* growth, independent of dexamethasone treatment. We used CRISPR/Cas9 to target *ikbkg* (inhibitor of nuclear factor kappa B kinase regulatory subunit gamma), which encodes IKKγ (NEMO) and is required for canonical NF-κB activation [20]. We designed two gRNAs targeting exons 2 and 3, both of which are part of the N-terminal IKKαβ binding domain (S7A Fig). We injected embryos at the 1 cell stage with both gRNAs targeting *ikbkg* or control gRNAs targeting *luciferase*, together with Cas9 protein. PCR using primers flanking the target sites on genomic DNA isolated from 2 dpf larvae confirmed successful targeting of DNA (S7B Fig). To test the survival of both control and macrophage-deficient larvae, we injected gRNAs and Cas9 into embryos from an *irf8^+/−^* in-cross and infected the resulting larvae with *A. fumigatus*. *irf8^+/+^ or irf8^+/−^ ikbkg* crispant larvae succumb to the infection at a significantly higher rate than larvae injected with control gRNAs (Fig 7A). In *irf8^−/−^* larvae, *ikbkg* targeting has an even larger effect on survival, demonstrating that when neutrophils are the primary immune cell type present, NF-κB activation plays a major role in promoting host survival (Fig 7A).

**Figure 7.**
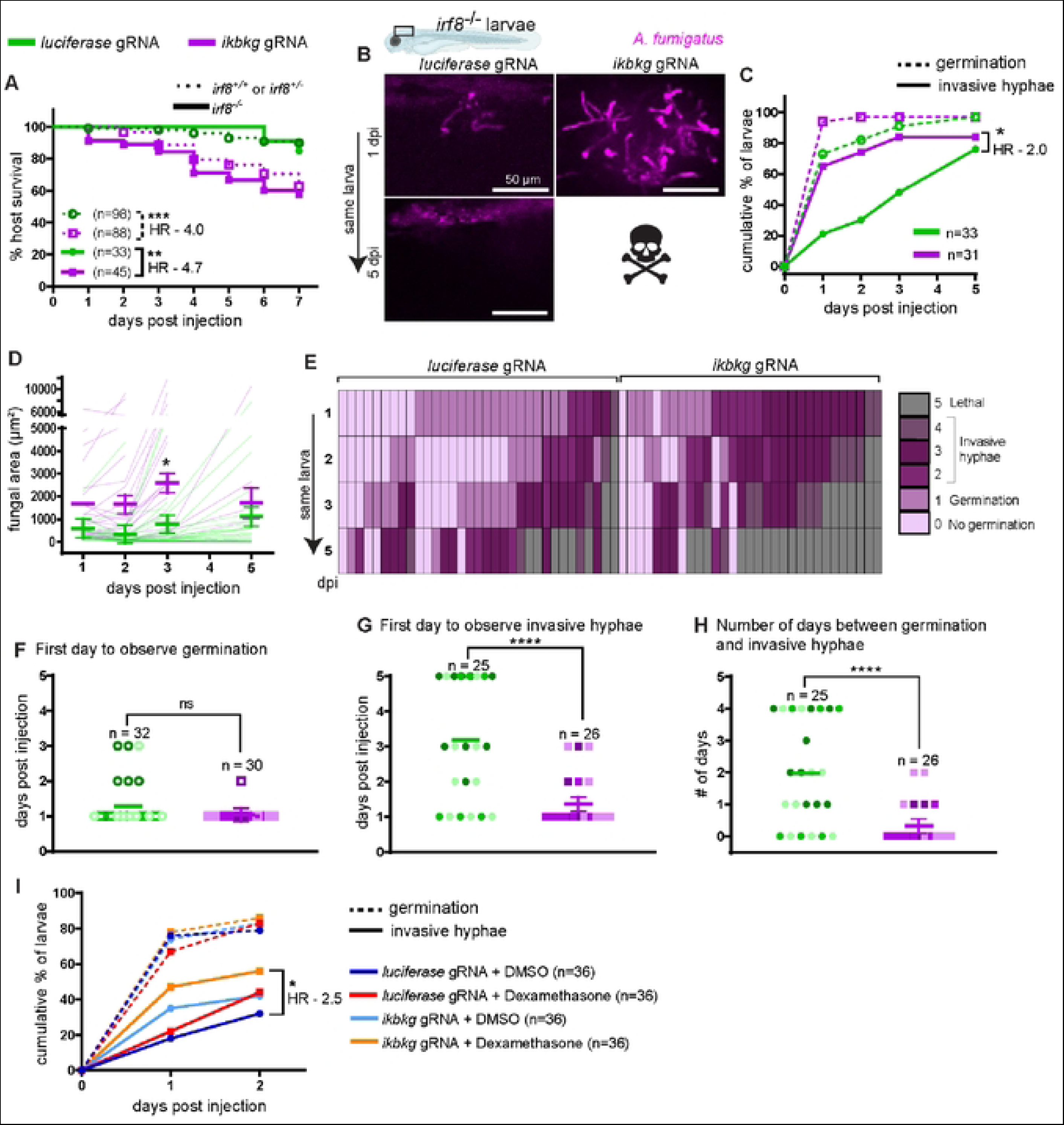
Neutrophils fail to control invasive hyphal growth in IKKγ crispant larvae. Macrophage-sufficient (*irf8^+/+^* or *irf8^+/−^*) and macrophage-deficient (*irf8^−/−^*) embryos at 1 cell stage were injected with Cas9 protein and 2 gRNAs targeting the IKKγ gene *ikbkg* or control gRNAs targeting *luciferase.* **(A)** Survival of larvae after injection with CEA10 spores at 2 dpf. Data are pooled from three independent replicates and the total larval N per condition is indicated. Cox proportional hazard regression analysis was used to calculate P values and hazard ratios (HR). Average injection CFUs: control + *luciferase* gRNA = 32, control + *ikbkg* gRNA = 30, *irf8^−/−^* + *luciferase* gRNA = 31, *irf8^−/−^* + *ikbkg* gRNA = 29. **(B-H)** *irf8^−/−^* embryos with labeled neutrophils (*Tg(lyz:BFP)*), injected with *ikbkg* or control gRNAs, were injected at 2 dpf with GFP-expressing TFYL49.1 spores and live imaged at 1, 2, 3, and 5 dpi. Data are pooled from three independent replicates, at least 10 larvae per condition, per replicate. (B) Representative images show hyphal growth differences in larvae injected with *ikbkg* or control gRNAs. Scale bar = 50 μm. (C) Cumulative percentage of larvae with germination (dotted line) and invasive hyphae (solid line) through 5 dpi. Cox proportional hazard regression analysis was used to calculate P values and hazard ratios (HR). (D) Fungal area was quantified from maximum intensity projection images. Bars represent emmeans ± SEM and P values were calculated by ANOVA. Each line represents an individual larva. (E) Severity of fungal growth was scored for all larvae and displayed as a heatmap. Representative images for each score can be found in S4 Fig. (F-H) In larvae in which fungal growth occurred, the day on which germination (F) and invasive hyphae (G) was first observed and the number of days between germination and invasive hyphae (H) are plotted. Bars represent emmeans ± SEM and P values calculated by ANOVA. Each data point represents an individual larva, color-coded by replicate. **(I)** *irf8^−/−^* larvae injected with *ikbkg* or control gRNAs were injected at 2 dpf with GFP-expressing TFYL49.1 spores, treated with 10 μM dexamethasone or DMSO, and live imaged at 1 and 2 dpi. The cumulative percentage of larvae with germination (dotted line) and invasive hyphae (solid line) through 2 dpi is shown. Data are pooled from three independent replicates, 12 larvae per condition, per replicate. Cox proportional hazard regression analysis was used to calculate P values and hazard ratios (HR).

To determine the requirement for NF-κB signaling in neutrophil-mediated control of invasive hyphae, we focused again on *irf8^−/−^* larvae and performed daily live imaging of *ikbkg* crispant larvae or control *luciferase-*targeted larvae after infection with a GFP-expressing strain of *A. fumigatus* (Fig 7B-H). As seen previously, almost all *irf8*^−/−^ larvae experience germination, regardless of gRNA injection (Fig 7A, B). However, the percentage of larvae with invasive hyphae is significantly higher in *ikbkg* crispant larvae compared to *luciferase-*targeted controls (Fig 7B, C). Additionally, *ikbkg* crispant larvae have higher fungal burdens as measured by fungal area, with statistically significantly higher burden at 3 dpi compared to control larvae (Fig 7D). Analysis of the severity of fungal growth indicates that control larvae are able to delay the development of invasive hyphal growth while *ikbkg* crispant larvae are not (Fig 7E). While germination appears at 1 dpi on average in both *ikbkg* crispant and control larvae (Fig 7F), invasive hyphae also appear as early as 1 dpi in *ikbkg* crispant larvae (Fig 7G). In control larvae, *A. fumigatus* takes ~2 days to develop invasive hyphae after germination while this occurs on the same day in *ikbkg* crispant larvae (Fig 7G, H). Together, these data demonstrate that in *irf8* mutant larvae in which neutrophils are solely responsible for fungal control, genetic NF-κB pathway inhibition through *ikbkg* (IKKγ) targeting with CRISPR/Cas9 phenocopies dexamethasone treatment. The effect of *ikbkg* targeting on neutrophil recruitment to the infection site is also similar to effect of dexamethasone (S8 Fig). These data are consistent with the conclusion that the major signaling pathway inhibited by dexamethasone to inhibit neutrophil-mediated control of invasive hyphal growth is NF-κB.

To test if dexamethasone treatment has other NF-κB independent effects, we exposed infected *irf8^−/−^*, *ikbkg* crispant or control *luciferase*-targeted larvae to dexamethasone or vehicle control. We then monitored fungal germination and hyphal growth by live imaging at 1 and 2 dpi. As expected, *A. fumigatus* germinates readily in larvae in all four conditions (Fig 7I). However, we did observe minor differences in the percentage of larvae harboring invasive hyphal growth. Control *luciferase* gRNA + DMSO larvae have the lowest rate of hyphal growth, while either dexamethasone treatment or *ikbkg* targeting alone increases this rate (Fig 7I). The combination of both *ikbkg* gRNA and dexamethasone exposure increases the percentage of larvae with invasive hyphae further (Fig 7I). These data suggest that dexamethasone may inhibit other pathways besides NF-κB that have minor roles in promoting neutrophil control of invasive hyphal growth. Overall, however, our data demonstrate that the primary effect of dexamethasone in causing host susceptibility to *A. fumigatus* infection is through inhibition of NF-κB-activated neutrophil functions against invasive fungal growth.

## Discussion

Prolonged glucocorticoid therapy is one of the major risk factors for invasive aspergillosis [7, 46]. The anti-inflammatory and immunosuppressive effects of glucocorticoids are largely caused by inhibition of NF-κB transcription factor activation and signaling [1]. *A. fumigatus* infection activates NF-κB *in vitro* and *in vivo* [26, 47], but it has been unclear if NF-κB suppression is the main mechanism of glucocorticoid-mediated susceptibility to invasive aspergillosis in these patients. Here, we used an *A. fumigatus*-zebrafish larvae infection system to demonstrate that glucocorticoids induce susceptibility in infected larvae through glucocorticoid receptor-mediated suppression of NF-κB. To elucidate this pathway, we used CRISPR/Cas9 to target the glucocorticoid receptor and the NF-κB pathway activator IKKγ. Glucocorticoid receptor and IKKγ are essential genes during development and cannot be easily targeted for infection studies in mice as mice lacking these genes are not viable [48, 49]. By using F0 mosaic crispant zebrafish larvae, we were able to inhibit the function of these genes in viable, morphologically normal, developing organisms. We find similar disease phenotypes in larvae exposed to dexamethasone and in IKKγ crispant larvae, demonstrating that inhibition of NF-κB is the major mechanism responsible for glucocorticoid-mediated susceptibility to *A. fumigatus* infection. However, IKKγ crispant larvae have a more severe disease phenotype when combined with dexamethasone, compared to either IKKγ mutation or dexamethasone alone, suggesting that dexamethasone also suppresses other minor pathways that contribute to control of fungal growth. One possible pathway that remains to be tested is activator protein-1 (AP-1) activation, which can be suppressed by glucocorticoids [50], and can be activated by *A. fumigatus* hyphae stimulation of mouse dendritic cells and alveolar macrophages [51, 52].

The key cellular immune mechanisms against *A. fumigatus* that are inhibited by glucocorticoid therapy have also been unclear. Glucocorticoids can affect all immune cell types to alter an inflammatory response [6]. Glucocorticoids potently suppress T-cell mediated responses, which are an integral component of anti-fungal immune mechanisms [6, 53]. However, larval zebrafish at the stage that was used for our study have not yet developed an adaptive system, and we have focused on the effects of dexamethasone on macrophages and neutrophils, cells that play key roles in host defense against different stages of fungal growth [26, 31, 32, 54]. Our results demonstrate that dexamethasone can still worsen *A. fumigatus* pathogenesis even before the involvement of the adaptive system [34]. To address whether dexamethasone primarily inhibits the function of macrophages or neutrophils or both, we used zebrafish larvae deficient for specific phagocyte populations. We find that dexamethasone does increase mortality in neutrophil-defective larvae, in which macrophages are the primary immune cells present, indicating that glucocorticoids can suppress macrophage-specific mechanisms. In monocytes and macrophages, glucocorticoids can suppress monocyte-to-macrophage maturation, chemotaxis to the infection site, phagocytosis, phagolysosomal fusion, oxidative killing, and pro-inflammatory cytokine secretion [6, 55]. In particular, in response to *Aspergillus* infections, glucocorticoids curb reactive oxygen species (ROS) production and autophagy-related protein recruitment to phagosomes downstream of ROS *in vitro* [31, 56]. Glucocorticoids can also alter macrophage M1 to M2 polarization, which can inhibit the ability of macrophages to prevent spore germination [57, 58]. In our study, we do find that dexamethasone treatment moderately inhibits macrophage recruitment to the infection site, but only at specific stages of infection, and even then, many macrophages are still present. We find no effect of dexamethasone on the ability of macrophages to kill spores or prevent germination *in vivo*. Additionally, the relative increase in susceptibility due to dexamethasone treatment in these larvae is actually less than in wild-type treated larvae, suggesting that macrophages are not the major target of this drug in inducing susceptibility to *A. fumigatus* infection.

In macrophage-deficient larvae, in which neutrophils are the primary immune cell present, dexamethasone increases mortality even more than in wild-type larvae, demonstrating that the inhibition of neutrophil function is a major target of dexamethasone-mediated susceptibility to *A. fumigatus* infection. In line with this conclusion, we report that dexamethasone curbs immune control of invasive hyphal growth, which is predominantly mediated by neutrophils. However, dexamethasone treatment does not prevent neutrophil migration to the infection site. We find large numbers of neutrophils in response to invasive hyphae in both wild-type and macrophage-deficient larvae in the presence of dexamethasone. Similarly, in rabbits treated with glucocorticoids, pulmonary lesions of invasive aspergillosis comprise neutrophil and monocyte infiltration and tissue necrosis [59]. Therefore, we conclude that dexamethasone inhibits the function of these cells against fungal infection. Neutrophils are efficient killers of hyphae and use multiple extracellular killing mechanisms such as degranulation, neutrophil extracellular traps (NETs), and the production of ROS [60]. Which of these mechanisms are most important against *A. fumigatus* and which are inhibited by dexamethasone treatment remains unclear. Neutrophils can undergo NETosis in response to *A. fumigatus* hyphae to control further growth [61–63], but the role of neutrophil-mediated ROS in *A. fumigatus* hyphal control is inconclusive [64, 65].

While our results demonstrate that dexamethasone inhibits NF-κB activation and this NF-κB activation promotes neutrophil functions against *A. fumigatus* hyphae, we cannot yet conclude whether this NF-κB activation is cell-intrinsic to neutrophils or non-cell-autonomous. We hypothesize that glucocorticoids inhibit activation of NF-κB in neutrophils, but cannot rule out the possibility that glucocorticoids inhibit activation of epithelial cells or macrophages, decreasing signals from these cells that activate neutrophils. Germination of spores exposes immunogenic ligands or pathogen-associated molecular patterns (PAMPs) such as β-glucans, which are otherwise masked by melanin and rodlet proteins in *A. fumigatus* spores, and this PAMP exposure induces a robust pro-inflammatory response that is likely mediated by all of these cells [66–68]. In our experiments, spore germination is the primary factor driving phagocyte recruitment, activation of an NF-κB reporter line, and induction of inflammatory cytokine expression, regardless of dexamethasone treatment. Our results demonstrate that excessive hyphal growth can override the suppression of NF-κB by dexamethasone.

Innate immune control of *A. fumigatus* varies with strain differences [26, 69]. In our experiments, we used spores derived from the CEA10 strain that was isolated from a patient [70]. CEA10-derived strains have relatively faster germination both *in vitro* and *in vivo* than spores of the other commonly studied AF293 strain [26, 71, 72]. As CEA10 is faster-germinating, resistance to infection with this strain in wild-type animals is more dependent on neutrophils and CEA10 is susceptible to neutrophil-mediated killing *in vitro* and *in vivo* [26, 32, 65], consistent with the primary activity of dexamethasone in causing susceptibility being inhibition of neutrophil functions against invasive hyphal growth. It is currently unclear if the role of glucocorticoids in the pathogenesis of more slowly germinating strains, like AF293, is the same. AF293 spores can reside within macrophage clusters, which inhibit their germination, and therefore evade neutrophil-mediated killing and persist in the host for long time [26]. Therefore, it could be that in these infections, glucocorticoid-mediated inhibition of macrophage function plays a larger role.

Glucocorticoid therapy is widely used in patients to decrease morbidity and mortality due to a variety of causes including transplant rejection and autoimmune disorders. However, a by-product of this therapy is increased susceptibility to infectious organisms, including fungi like *Aspergillus* spp. In order to better treat these opportunistic infections, it is important to fully understand the molecular and cellular mechanisms that are inhibited by glucocorticoid therapy that are the proximate cause of susceptibility to infection. Here, using a larval zebrafish host, we find that the major innate immune cell type inhibited by this therapy is neutrophils and that this inhibition allows for uncontrolled invasive hyphal growth in infected animals. However, the specific functions of neutrophils that are inhibited remain to be uncovered. In this model, NF-κB is the major molecular pathway inhibited by glucocorticoid treatment. However, the role of NF-κB in human infection is unclear, as the risk of invasive aspergillosis did not depend on single nucleotide polymorphisms (SNPs) of NF-κB and NF-κB pathway components in a cohort of hematopoietic stem-cell transplant patients [73], and the effect of glucocorticoids on other pathways should also be an area of future study.

## Materials and Methods

### Ethics statement

All experimental procedures of zebrafish embryos and larvae were performed, and adult zebrafish were maintained and handled, according to protocols approved by the Clemson University Institutional Animal Care and Use Committee (AUP2021-0109, AUP2022-0093, AUP2022-0111). Larvae were anesthetized using buffered tricaine prior to any experimental procedures. Larvae were euthanized at 4°C and adults were euthanized with buffered tricaine.

### Zebrafish lines and maintenance

Adult zebrafish were maintained at 28°C at 14/10 hr light/dark cycles. All mutant and transgenic lines were maintained in the wild-type AB background, and are listed in Table 1. Embryos were collected after natural spawning, and were maintained in E3 medium with methylene blue at 28°C. Embryos were manually dechorionated and anesthetized in 0.3 mg/mL buffered tricaine prior to any experimental procedures. Larvae used for imaging were treated with 200 μM N-phenylthiourea (PTU) starting from 24 hours post fertilization to inhibit pigment formation. All transgenic larvae were screened for fluorescent expression prior to infections. The *irf8* mutant adults were maintained as *irf8^+/−^* with additional transgenes expressing fluorescent markers in neutrophils (*Tg(mpx:mCherry)* or *Tg(lyzC:BFP)*). Larvae from *irf8^+/−^* in-crosses were screened for a high number of neutrophils to select for *irf8^−/−^* individuals prior to infections and larvae were genotyped after the experiments were concluded as previously described [45], using the primers F: 5’ CAGGAGAGTTCAGTAAATTGAGC 3’; R: 5’ CTTGTTTTCCCGCATGTTTCC 3’.

**Table 1.**
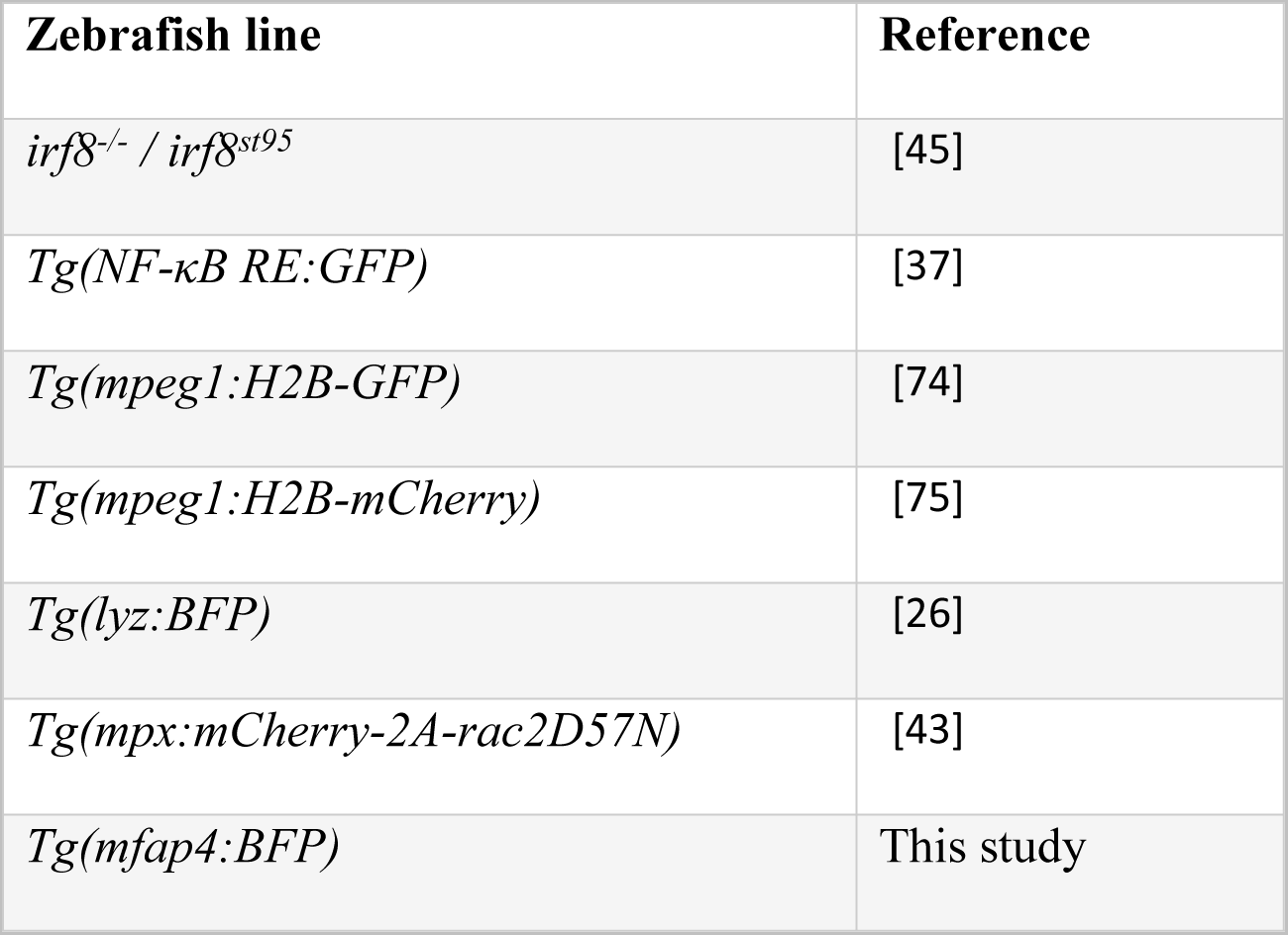
Zebrafish lines used in this study.

| Zebrafish line | Reference |
| --- | --- |
| *irf8*<sup>-/-</sup> / *irf8*<sup>st95</sup> | [45] |
| *Tg(NF-κB RE:GFP)* | [37] |
| *Tg(mpeg1:H2B-GFP)* | [74] |
| *Tg(mpeg1:H2B-mCherry)* | [75] |
| *Tg(lyz:BFP)* | [26] |
| *Tg(mpx:mCherry-2A-rac2D57N)* | [43] |
| *Tg(mfap4:BFP)* | This study |

### Generation of *Tg(mfap4:BFP)* fish line

First, a clean Tol2 vector containing just the *mfap4* promoter was generated. Tol2-*mpx:mCherry-2A-rac2* [43] (a gift from Anna Huttenlocher) was cut with NheI and SalI (Promega) to remove the *mpx:mCherry-2A-rac2* insert. The *mfap4* promoter was then amplified from p5E-*mfap4* (Addgene #70052, a gift from David Tobin) for InFusion cloning (Takara Bio) into the digested Tol2 backbone (F: 5’ GAAGTAAAAGGCTAGC*GCGTTTCTTGGTACAGCTG* 3’; R: 5’ TTCTAGATCAGTCGAC*CACGATCTAAAGTCATGAAG* 3’). To generate Tol2-*mfap4:BFP*, *BFP* was amplified from Tol2-*lyz:BFP* [26] (a gift from Anna Huttenlocher) for HiFi cloning (NEB) into the Tol2-*mfap4* vector cut open with SalI (F: 5’ TGACTTTAGATCGTGGTCGAC*GGTACCTCGCCACCATGA* 3’; R: 5’ CTATAGTTCTAGATCATCGAC*TCACTTGTGCCCCAGTTT* 3’).

For integration of *mfap4:BFP* into the zebrafish genome, *Tol2 transposase* was *in vitro* transcribed from NotI-digested (NEB) pCS2-*transposase* (a gift from Anna Huttenlocher) using an mMESSAGE mMACHINE SP6 kit according to the manufacturer’s directions (Invitrogen). mRNA was purified with a MEGAclear kit (Invitrogen). 1-3 nl of an injection mix containing 20 ng/µl Tol2-*mfap4:BFP* plasmid and 10 ng/µl *transposase* mRNA was injected into the yolk of single cell embryos of the AB strain. Injected F0 embryos were grown to adulthood and a founder with integration of the DNA into the germline was determined by outcrossing single F0 adults and screening for BFP expression.

### *Aspergillus fumigatus* strains and spore preparation for injections

The CEA10 strain [70] and a CEA10-derived GFP-expressing TFYL49.1 strain [76] were used for non-imaging and imaging experiments, respectively. These strains are equivalent in pathogenesis in zebrafish larvae [26]. Spores were prepared as previously described [27]. Briefly, 10^6^ spores were spread on 10 cm plates with solid glucose minimal media (GMM) and were incubated at 37°C for 4 days. Spores were collected into sterile water with 0.01% Tween by scraping using a L spreader and were filtered through sterile miracloth (Sigma-Aldrich) into a 50 mL centrifuge tube. Spores were pelleted by centrifugation at 900 g for 10 mins, washed in sterile PBS, pelleted again, and finally resuspended in 5 mL of sterile PBS. This suspension was again filtered through miracloth into a new 50 mL tube. The spore concentration was determined using a hemocytometer and a suspension at 1.5 × 10^8^/ mL was made in PBS and stored at 4°C for up to ~6 weeks.

### *In vitro* fungal growth assay

10 cm solid GMM plates containing 20 mL GMM agar with 10 μM dexamethasone or 0.01% DMSO vehicle control were prepared and stored at 4°C for up to ~4 months. Spores of the GFP-expressing TFYL49.1 strain were prepared as described above and resuspended at 10^7^/ mL and stored at 4°C for up to ~6 weeks. 2 μL was dispensed into the middle of the plate and plates were incubated at 37°C for 4 days. To quantify the growth in each condition, the diameter of the colony was measured daily. To quantify branching of the growing hyphae, at 2 days post culture, a piece of agar from the edge of each colony was cut out and placed on a glass slide and flattened using a coverslip. Hyphae in this piece were imaged using a confocal microscope, as described below. This experiment was repeated twice with three plates/condition/experiment. One slide per plate was used for imaging, and four microscopic fields were captured for each slide.

### Live-dead spore labeling

Spores of the GFP-expressing TFYL49.1 strain were stained with AlexaFluor546 as described previously [39, 77, 78]. Briefly, spores were incubated on a shaker with 0.5 mg/mL of biotin-XX, SSE (Molecular Probes) in 0.05M NaHCO_3_ at 4°C for 2 hrs. Spores were pelleted by centrifugation and washed twice with 100 mM Tris-HCl (pH 8.0) on a shaker at 4°C for 30 min to deactivate free-floating biotin. Spores were washed with 1X PBS twice. Spores were then resuspended in 1X PBS containing 20 μg/mL of streptavidin-AlexaFluor546 (Invitrogen) and incubated for 40 min at room temperature. Stained spores were then pelleted and resuspended in 1X PBS, spore concentration was enumerated, and a spore suspension was made at 1.5 × 10^8^/ mL and stored at 4°C for up to ~4 weeks.

### Zebrafish hindbrain microinjections

*A. fumigatus* spores were injected into the hindbrain ventricle of 2 days post fertilization (dpf) larvae as described previously [36]. The prepared 1.5 × 10^8^/ mL spore suspension was mixed at 2:1 with filter-sterilized phenol red to achieve a final concentration of 1 × 10^8^/ mL. Injection plates made of 2% agarose in E3 were coated with 2% filter-sterilized bovine serum albumin (BSA) to prevent larvae sticking to the agarose. Dechorionated and anesthetized 2 dpf larvae were transferred to and aligned on the injection plate. A microinjection setup supplied with pressure injector, micromanupulator, micropipet holder, footswitch, and back pressure unit (Applied Scientific Instrumentation) was used to inject 30-50 spores into individual larvae. Actual injection doses were monitored by CFU plating as described below and are reported in each figure legend. PBS mixed with phenol red was used as a mock infection control. Injected larvae were then rinsed at least twice with E3 without methylene blue to remove tricaine and any free spores. For imaging experiments, larvae were returned to E3 containing 200 µM PTU. Larvae were transferred to 96-well plates for survival and CFU experiments, 48-well plates for daily imaging experiments, or 6-well plates for RNA extraction and single day imaging experiments.

### Morpholino injections

A stock solution of *pu.1* (*spi1b*) morpholino oligonucleotide (MO) (ZFIN MO1-*spi1b*: 5’ GATATACTGATACTCCATTGGTGGT 3’) (GeneTools) was made by resuspension in water to 1 mM and stored at 4°C [41, 79]. For injections, the stock was diluted in water with 0.5X CutSmart Buffer (New England Biolabs) and 0.1% filter-sterilized phenol red. We used 2 different concentrations of *pu.1* MO: low-dose 0.05 mM to prevent development of macrophages [44] or high-dose 0.5 mM to prevent development of both macrophages and neutrophils [41]. Optimization of low-dose *pu.1* MO to only inhibit macrophage development is mentioned below. A standard control MO at matching concentrations was used as a control. 3 nL of the injection mix was injected into the yolk of 1-2 cell stage embryos. Efficacy of 0.5 mM *pu.1* knockdown was determined by injection into embryos of macrophage-(*Tg(mpeg1:H2B-GFP)*) and neutrophil-labeled (*Tg(mpx:mCherry)*) zebrafish lines and larvae were screened for the loss of fluorescent signal using a fluorescent zoomscope (Zeiss SteREO Discovery.V12 PentaFluar with Achromat S 1.0x objective) prior to *A. fumigatus* infections. To identify a low concentration of *pu.1* MO that only inhibits macrophage development but not neutrophil development, multiple different concentrations ranging from 0.05 mM to 0.5 mM were tested. Embryos of macrophage-labeled (*Tg(mpeg1:H2B-GFP)*) and neutrophil-labeled (*Tg(mpx:mCherry)*) zebrafish lines were injected with increasing concentrations of *pu.1* MO and larvae at 2 dpf were screened for the loss of GFP signal but intact mCherry signal using a fluorescent zoomscope (Zeiss SteREO Discovery.V12 PentaFluar with Achromat S 1.0x objective). To further test if neutrophils are still active with low-dose *pu.1* MO injections, we performed a tail wounding experiment as previously described [80]. Briefly, the tails of the larvae injected with 0.05 mM *pu.1* or control MO were transected using a no.10 Feather surgical blade (GF Health Products) and the larvae were confocal imaged (as described in the Live Imaging section) at 2 hours post wounding. The number of neutrophils at the wounding site was enumerated. 6 larvae were used for each condition, and the experiment was done once.

### CRISPR gRNA design and injections

For each target gene, two guide RNAs (gRNA) were designed to bind to regions of the coding sequence that are required for the function of the protein and in which the translated amino acid sequence is conserved with the human protein. gRNA targets were identified with the CHOPCHOP web-based program [81–83]. gRNA sequences are listed in Table 2. To generate DNA templates for *in vitro* transcription of gRNAs, gene-specific oligo sequences were generated containing a T7 promoter (5’-TAATACGACTCACTATAG-3’), the target sequence, and an overlap region to pair with a constant oligo encoding the reverse complement of the Cas9 binding sequence (Integrated DNA Technologies)(constant oligo (5’-AAAAGCACCGACTCGGTGCCACTTTTTCAAGTTGATAACGGACTAGCCTTATTTTAA CTTGC TATTTCTAGCTCTAAAAC-3’)). Gene-specific and constant oligos were annealed and T4 DNA polymerase (New England Biolabs) was added to generate the DNA template. Purified template was *in vitro* transcribed using T7 RNA polymerase (New England Biolabs), treated with DNase I (New England Biolabs), and purified using a Monarch RNA cleanup kit (New England Biolabs). Embryos were injected with both gRNAs targeting a single gene. An injection mix containing 75 ng/μL of each gRNA and 250 ng/μL Cas9 protein (PNA Bio) was used. Two control gRNAs targeting *luciferase (luc)* at matching concentrations were used as the control. 1-2 nL of injection mix was injected into the yolk of 1-cell stage embryos. Genomic DNA from individual larvae was extracted at 2 dpf in 50 mM NaOH at 95°C. The efficacy of gRNAs were tested by PCR using primers flanking the target sequence. We used two sets of PCR, one with primers flanking an individual target site (F1R1 or F2R2) and with primers flanking the two target sequences (F1R2) as shown in S2 and S7 Figs. For *ikbkg*1+2-injected larvae, a separate primer pair flanking the two target sites with matching Tm was designed (F3R3). The primers used are listed in Table 3. The PCR products were run on a 2.5% agarose gel to visualize alterations of genomic DNA as shown in S2 and S7 Figs.

**Table 2.**
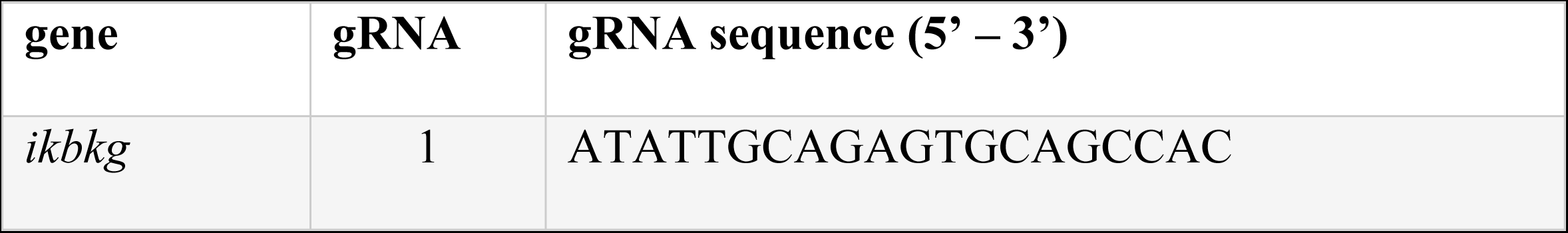
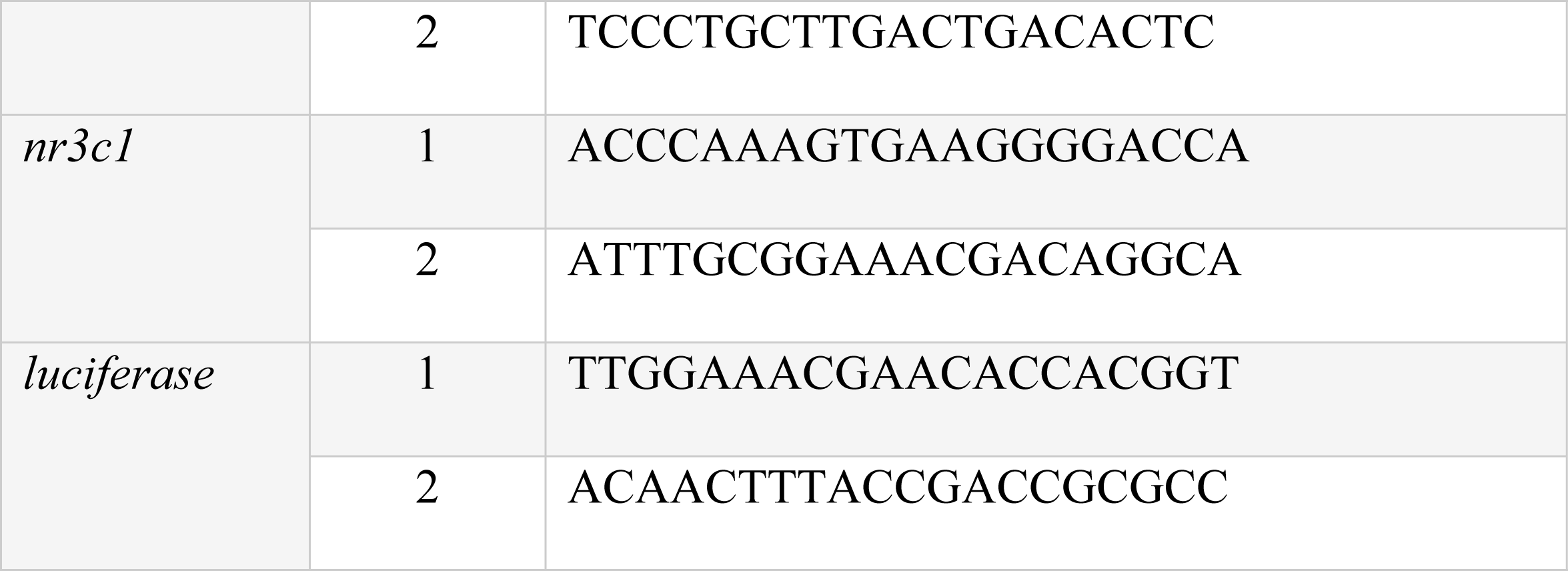
CRISPR gRNAs.

**Table 3.**
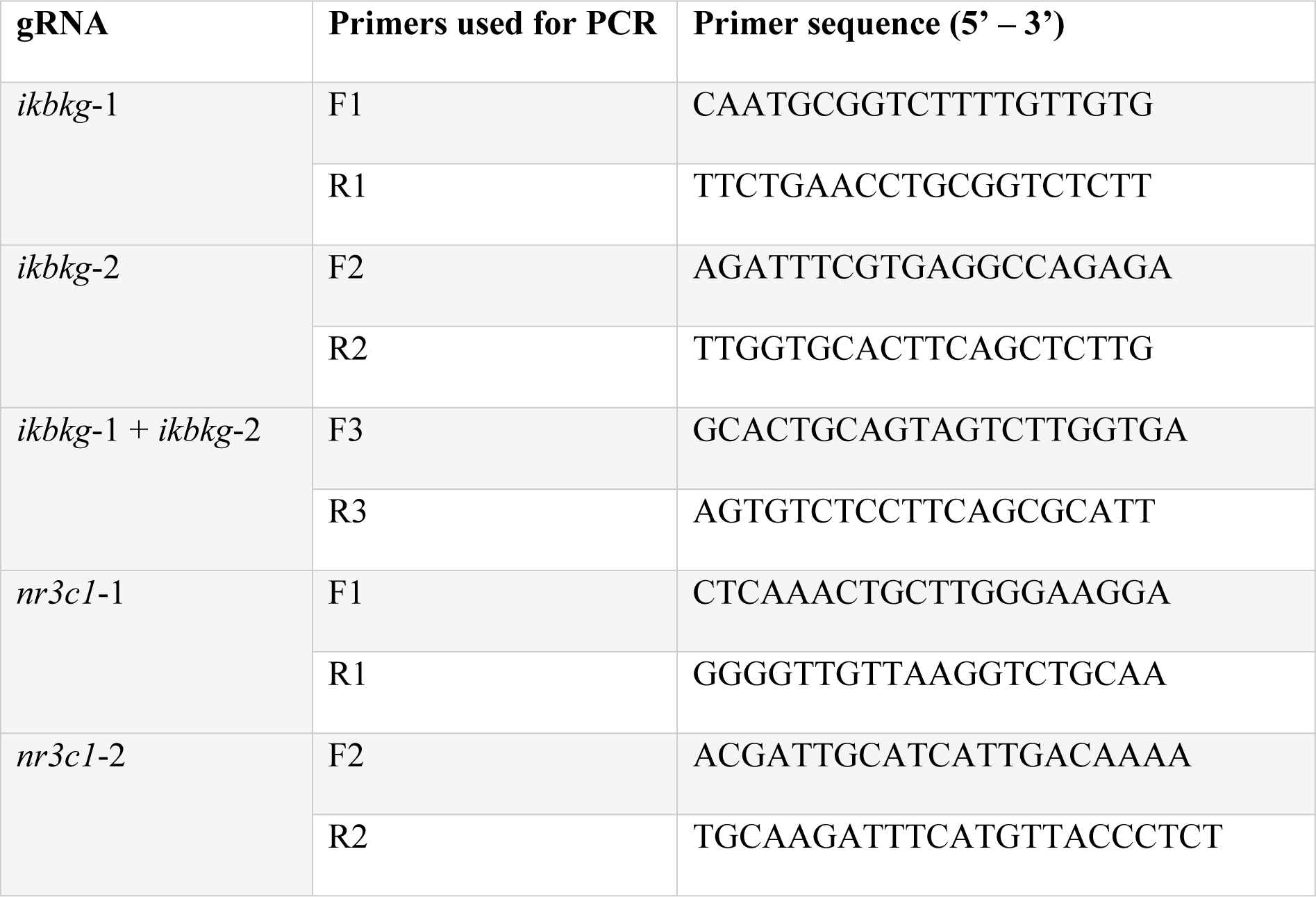
Primers used to test successful alteration of DNA.

### Clodronate liposome injections

Larvae expressing GFP in macrophages (*Tg(mpeg1:H2B-GFP)*) were manually dechorionated at 1 dpf. Clodronate or PBS liposomes (Liposoma) were stored at 4°C. Prior to injections, 10% of volume of filter-sterilized phenol red was added to the liposomes and 2 nL was i.v. injected into the caudal vein plexus of larvae. To confirm macrophage depletion, larvae were screened for the loss of GFP signal using a fluorescent zoomscope (Zeiss SteREO Discovery.V12 PentaFluar with Achromat S 1.0x objective) prior to *A. fumigatus* infection at 2 dpf.

### Drug treatments

Infected larvae were exposed to dexamethasone (Sigma-Aldrich) at 10 μM, a concentration which was previously used in zebrafish larvae [26]. A 1000X 10 mM stock was prepared in DMSO and 0.1% DMSO was used as the vehicle control. Directly after injection, E3 was removed from dishes containing larvae and new E3 with pre-mixed dexamethasone or DMSO was added. Larvae were kept in the same solution for the entirety of the experiment. For daily imaging experiments, larvae were pipetted out of the drug treatment, imaged, and were transferred back into the same drug solution.

### CFU counts

Single larvae were transferred to 1.7 mL microcentrifuge tubes into 90 μL of PBS containing 1 mg/mL ampicillin and 0.5 mg/mL of kanamycin. Larvae were euthanized at 4°C overnight and homogenized with a tissue lyser (Qiagen) at 1800 oscillations/min (30 Hz) for 6 mins. The suspension was then centrifuged at 17000 g for 30 seconds, resuspended by pipetting, and spread on a GMM plate. Plates were incubated at 37°C for 3 days and the number of colonies were counted. For survival experiments, 8 larvae for each condition were collected and euthanized immediately after injections, and were plated the next day to enumerate the actual injection dose. To monitor the fungal burden across multiple days, 8 larvae/condition/day were plated and CFU counts were normalized to the initial injection dose for each condition.

### RNA extraction and RT-qPCR

To quantify cytokine gene expression, larvae were infected with TFYL49.1 spores and exposed to dexamethasone or DMSO. At 1 or 2 dpi, infected larvae were anesthetized and screened using a Zeiss Cell Observer Spinning Disk confocal microscope to split larvae into groups based on the presence of germinated spores. From this screening, the pooled no germination group contained 20 larvae and the pooled germination group contained 1-3 larvae per replicate. 500 ng of RNA was used for cDNA synthesis. To quantify *ikbkb* expression in wild-type larvae, larvae were injected with CEA10 spores or PBS and were exposed to dexamethasone or DMSO. 20 pooled larvae for each condition per replicate at 1 or 2 dpi were used for RNA extraction. 1000 ng of RNA was used for cDNA synthesis. To test *ikbkb* expression in glucocorticoid receptor targeted larvae, *nr3c1* crispant or control larvae were exposed to dexamethasone or DMSO at 2 dpf. At 1 day post treatment (dpt), 20 pooled larvae from each condition per replicate were used for RNA extraction. 1000 ng of RNA was used for cDNA synthesis.

For RNA extraction, anesthetized larvae were transferred to a 1.7 mL microcentrifuge tube homogenized in 500 μL TRIzol (Invitrogen) on a disruptor genie for 10 minutes. RNA was extracted following the manufacturer’s instructions, using 4 μg of glycogen as a carrier. cDNA synthesis was done with iScript RT Supermix with oligo dT (Bio-Rad). cDNA was diluted 1:10 and 4 μL of diluted cDNA was used for qPCR in a 10 μL reaction, using SYBR Green Supermix (Bio-Rad) and primers listed in S1 Table. Fold change was calculated with the ΔΔC_q_ method, using *rps11* as the house-keeping gene [84] and three independent replicates were performed for each RT-qPCR.

### Live imaging

Fluorescent positive larvae were screened on using a fluorescent zoomscope (Zeiss SteREO Discovery.V12 PentaFluar with Achromat S 1.0x objective) prior to *A. fumigatus* infections. Larvae were imaged using a Zeiss Cell Observer Spinning Disk confocal microscope on a Axio Observer 7 microscope stand with a confocal scanhead (Yokogawa CSU-X) and a Photometrics Evolve 512 EMCCD camera. A Plan-Apochromat 20X (0.8 NA) or an EC Plan-Neofluar 40X (0.75 NA) objective and ZEN software were used to acquire Z-stack images of the hindbrain area with 2.5 or 5 μm distance between slices. For daily imaging experiments, larvae were pipetted out of 48-well plates one at a time, anesthetized in tricaine, and transferred to a zWEDGI device [85, 86]. After imaging, larvae were rinsed in E3 with 200 µM PTU and transferred back into the original wells into the same drug solution. For single time point imaging, larvae were anesthetized in 6-well plates and were transferred to and imaged in a zWEDGI device [85, 86]. For experiments in the *irf8* mutant line, genomic DNA was isolated from whole larvae in 50 mM NaOH for genotyping when larvae were euthanized due to increased fungal growth or immediately after completion of imaging.

### Image analysis

All image analysis was performed blinded and with Image J/Fiji [87]. For any analysis where fluorescent intensity was quantified, images were not processed prior to analysis. To quantify the GFP signal from the *NF-κB RE:GFP* zebrafish line, the hindbrain area was manually identified using the polygon selection tool from the corresponding bright field image. The GFP signal was quantified within the identified area using maximum intensity projection of six z-slices containing *A. fumigatus* spores or hyphae. The displayed images show signal intensity with the 16 colors lookup table. To quantify phagocyte recruitment, images were processed with bilinear interpolation to increase pixel density two-fold prior to counting and the number of phagocytes and/or the phagocyte cluster area were quantified. Phagocyte numbers were manually counted across z-stacks using the Cell Counter plugin. Macrophage cluster area was measured in maximum intensity projections using the polygon selection tool. Displayed images were processed with bilinear interpolation to increase pixel density two-fold and maximum intensity projections of merged z-stacks were used. For live-dead staining, images were processed with bilinear interpolation to increase pixel density two-fold and live versus dead spores were counted using the Cell Counter plugin. The displayed images of live versus dead spores show a merged z-projection of 3 slices and were processed with gaussian blur (radius = 1) in Fiji to reduce noise. Fungal growth was manually categorized as germination or invasive hyphae. Any hyphal growth (branched or not) was considered an incidence of germination and the presence of branched hyphae was considered an incidence of invasive growth. 2D fungal area was measured by thresholding the fluorescent intensity from maximum intensity projections. To generate the heatmap of fungal growth, the severity was scored using pre-determined categories [39]. The scoring was: 0 – no germination, 1 – at least one event of germination, 2 – at least one event of branched hyphae, but hyphae restricted to the infection site, 3 – at least one event of branched hyphae, but hyphae spreading in the hindbrain ventricle, 4 – hyphae invading into nearby tissue, and 5 – lethal. Representative images of each category are shown in S4 Fig. Maximum intensity projection of z-stacks was used for the displayed images. The same experiment and the same images were used to enumerate the number of phagocytes and fungal growth. For *irf8^−/−^* larvae imaging experiments, neutrophil cluster area was measured from maximum intensity projections using the polygon selection tool. Maximum intensity projections of z-stacks were used for the displayed images. For the *in vitro* assay images, the fungal area was measured by quantifying GFP signal by thresholding the fluorescent intensity from the maximum intensity projection of all slices. The number of nodes/branching points were manually counted using the Cell Counter plugin.

### Statistical analysis

For all experiments, unless stated otherwise, pooled data from at least three independent replicates were generated and the total Ns are given in each figure. R version 4.1.0 was used for statistical analysis and GraphPad Prism version 10 (GraphPad Software) was used to generate graphs. Larval survival data and the cumulative percentage of larvae with fungal germination or invasive hyphae were analyzed by Cox proportional hazard regression to calculate P values and hazard ratios (HR). HR reports the likelihood of larvae succumbing to the infection in a particular condition as compared to the control. The statistical analysis considers variability within and between replicates to calculate the P values. Fluorescent intensity, phagocyte numbers and cluster area, fungal area, CFU counts, live-dead imaging analysis, and day of germination or invasive hyphae were analyzed with analysis of variance (ANOVA). For each condition, estimated marginal means (emmeans) and standard error (SEM) were calculated and pairwise comparisons were performed with Tukey’s adjustment. The statistical analysis considers variability within and between replicates to calculate the P values. The graphs for these analyses show values from individual larvae over time as individual lines or points in dot plots, and bars represent pooled emmeans ± SEM. The points in dot plots are color-coded by replicate. For RT-qPCR, the fold change was analyzed by t-test in Excel. For *in vitro* data, the radius of the colonies and the number of nodes normalized to the fungal area were compared by t-test in Excel. In the dot plot, each dot represents an individual plate, and the dots are color-coded by replicate.

## Acknowledgements

We thank Celia Shiau for providing the *irf8* mutant zebrafish line. We thank Anna Huttenlocher for sharing all other transgenic lines. We thank Sourabh Dhingra for helping to design the *A. fumigatus in vitro* assay. We thank all the members of the Rosowski Lab for helpful discussions.

## Supporting Information

**S1 Fig. Dexamethasone suppresses NF-κB activation. (A)** Larvae of NF-κB reporter line (*Tg(NF-κB RE:GFP)*) were injected with CEA10 spores and were exposed to 10 μM dexamethasone or DMSO vehicle control. Fluorescent expression was quantified in the hindbrain ventricle from z projections at 2 dpi. Quantification data are shown with emmeans ± SEM from three independent replicates and the total larval N per condition is indicated. Each data point represents an individual larvae, color-coded by replicate. P values were calculated by ANOVA. **(B)** Larvae were injected with GFP-expressing TFYL49.1 (CEA10) spores and exposed to 10 μM dexamethasone or DMSO. At 1 and 2 dpi, larvae were screened for germination and total RNA was extracted from each pooled group. RT-qPCR analysis of cytokine expression in larvae is shown. Data are normalized to DMSO no germination control group. P values were calculated by Student’s t-test. Data are from three independent replicates.

**S2 Fig. Generation of glucocorticoid receptor crispant larvae. (A-B)** Design and efficiency of *nr3c1* gRNAs. (A) Zebrafish glucocorticoid receptor (*nr3c1*) gene structure and the target sites for the two gRNAs and primers used for PCR. (B) 1 cell stage embryos were injected with gRNAs targeting *nr3c1* or control gRNAs targeting *luciferase* together with Cas9 protein. At 2 dpf, genomic DNA was extracted from individual larvae. Successful targeting of DNA was confirmed by PCR using primer pairs illustrated in (A). Gel electrophoresis indicates clean bands for control larvae with F1R1 and F2R2 primers, while for the gRNA-injected larvae, the bands are blurry indicating random mosaic insertions and deletions. F1R2 primer pair indicates that ~36 kb piece of DNA can be excised in larvae injected with both target gRNAs. No PCR band is detected in control larvae due to the large amplicon size. **(C)** Survival of *nr3c1* mutant or control larvae injected with PBS and exposed to 10 μM dexamethasone or DMSO. Data are pooled from three independent replicates, at least 22 larvae per condition, per replicate and the total larval N per condition is indicated in each figure. Cox proportional hazard regression analysis was used to calculate P values and hazard ratios (HR).

**S3. Fig. Dexamethasone does not affect *A. fumigatus* spore germination *in vitro*.** GFP-expressing TFYL49.1 (CEA10) spores were inoculated into the middle of solid GMM plates containing 10 μM dexamethasone or DMSO vehicle control. **(A)** The diameter of the colony was measured after 1-4 days. **(B-C)** At 2 days post culture, a sample from the edge of the colony of each plate was transferred to a glass slide and imaged with a confocal microscope. Four fields were imaged for each slide. Data are pooled from two independent replicates, and three plates per condition, per replicate were used. (B) A representative image showing branched hyphae; yellow arrows point to nodes. Scale bar = 100 μm. (C) Per field of view, the number of nodes were counted and normalized to the total fungal area. Fungal area was quantified from maximum intensity projection images. Bars represent means ± SEM and P values calculated by Student’s T-test. Each data point represents the average value from individual plates, color-coded by replicate.

**S4 Fig. Representative images of categories of *A. fumigatus* hyphal growth.** Wild-type larvae were injected with GFP-expressing TFYL49.1 (CEA10) spores at 2 dpf, exposed to 10 μM dexamethasone or DMSO vehicle control and live imaged at 1, 2, 3, and 5 dpi. Incidences of hyphal growth were scored depending on the extent of hyphal growth. Category 1: presence of a germ tube. Category 2: presence of branched hyphae (filled arrow), yet small fungal bolus. Category 3: presence of spread-out invasive hyphae. Category 4: presence of severe invasive hyphae and tissue damage. Scale bar: 50 μm or 25 μm.

**S5 Fig. Survival of phagocyte-deficient larvae and validation of macrophage depletion with low concentration *pu.1* morpholino.** (A-D) Survival of larvae injected at 2 dpf with CEA10 *A. fumigatus* spores or PBS mock-infection in the presence of 10 μM dexamethasone or DMSO vehicle control. (A) Survival of phagocyte-depleted or control larvae injected with CEA10 spores. Development of all phagocytes was inhibited by 0.5 mM of *pu.1* morpholino (MO). Control larvae received standard control MO. Data are pooled from two independent replicates, at least 23 larvae per condition, per replicate and the total larval N per condition is indicated in the figure. Average injection CFUs: control MO = 55, *pu.1* MO = 65. (B) Survival of neutrophil-defective larvae (*mpx*:*rac2D57N*) and wild-type larvae after PBS-mock infection. (C, D) Macrophages were depleted via clodronate liposome i.v. injection at 1 dpf. Control larvae received PBS liposomes. Survival of macrophage-depleted larvae injected with (C) CEA10 spored or (D) PBS mock-infection. Average injection CFUs: PBS liposomes = 23, clodronate liposomes = 20. **(E-H)** Development of macrophages was inhibited by 0.05 mM *pu.1* MO. Control larvae received standard control MO. (E) Low dose *pu.1* MO or control larvae expressing fluorescent markers in macrophages (*Tg(mpeg1:H2B-GFP)*) or neutrophils (*Tg(mpx:mCherry)*) were imaged around the caudal hematopoietic area to visualize phagocytes at 2 dpf. Representative images show a lack of macrophages but intact neutrophils. Scale bar = 100 μm. (F, G) Low dose *pu.1* MO or control larvae with fluorescent neutrophils (*Tg(mpx:mCherry)*) were wounded by tail transection and were imaged at 2 hours post injury (hpi). (F) Representative images showing neutrophil recruitment to the wounding site. Scale bar = 50 μm. (G) Quantification of the number of neutrophils at the wound site is shown with means ± SEM from one replicate and the total larval N per condition is indicated. Each data point represents an individual larva. (H) Survival of low dose *pu.1* MO macrophage-deficient or control larvae after PBS mock-infection is shown. (B-D, H) Data are pooled from three independent replicates, at least 9 larvae per condition, per replicate and the total larval N per condition is indicated in each figure. Cox proportional hazard regression analysis was used to calculate P values and hazard ratios (HR).

**S6 Fig. Dexamethasone does not significantly affect neutrophil recruitment to the infection site in *irf8^−/−^* larvae.** *irf8^−/−^* larvae with labeled neutrophils (*Tg(lyz:BFP)*) were injected with GFP-expressing TFYL49.1 (CEA10) spores at 2 dpf, exposed to 10 μM dexamethasone or DMSO vehicle control and live imaged at 1, 2, 3, and 5 dpi. Data are pooled from three independent replicates, at least 10 larvae per condition, per replicate. **(A)** Fungal area was quantified from maximum intensity projection images. **(B)** Neutrophil cluster area was quantified from maximum intensity projection images. (A, B) Bars represent emmeans ± SEM and P values were calculated by ANOVA. Each line represents an individual larva. **(C)** Neutrophil cluster area one day before germination occurred, on the day of germination, and on the day invasive hyphae occurred were quantified for larvae that experienced fungal growth. Bars represent emmeans ± SEM and P values were calculated by ANOVA. Each data point represents an individual larva, color-coded by replicate. **(D)** The correlation of neutrophil cluster area to fungal area was plotted for larvae that had germination and invasive hyphae. All larvae have had neutrophil clusters at some point during infection.

**S7 Fig. Generation and validation of IKKγ-deficient *ikbkg* crispant larvae. (A) Zebrafish** IKKγ (*ikbkg*) gene structure and the target sites for gRNAs and primers used for PCR. **(B)** 1 cell stage embryos were injected with gRNAs targeting *nr3c1* or control gRNAs targeting *luciferase* together with Cas9 protein. At 2 dpf, genomic DNA was extracted from individual larvae. Successful targeting of DNA was demonstrated by PCR using primer pairs in (A). Gel electrophoresis indicates clean bands for control larvae with F1R1 and F2R2 primers, while for the target gRNA-injected larvae, the bands are blurry indicating random mosaic insertions and deletions. F1R2 primer pair indicates that ~1.6 kb piece of DNA can be excised in larvae injected with both target gRNAs. No PCR band is detected in control larvae due to the large amplicon size.

**S8 Fig. IKKγ targeting does not significantly inhibit neutrophil recruitment to *A. fumigatus* infection.** *irf8^−/−^* embryos with labeled neutrophils (*Tg(lyz:BFP)*) were injected with *ikbkg* or control gRNAs. At 2 dpf, larvae were injected with GFP-expressing TFYL49.1 spores and live imaged at 1, 2, 3, and 5 dpi. Data are pooled from three independent replicates, at least 10 larvae per condition, per replicate. **(A)** Neutrophil cluster area was quantified from maximum intensity projection images. Bars represent emmeans ± SEM and P values were calculated by ANOVA. Each line represents an individual larva. **(B)** Neutrophil cluster area one day before germination occurred, on the day of germination, and on the day invasive hyphae occurred were plotted for larvae that experienced fungal growth. Bars represent emmeans ± SEM and P values were calculated by ANOVA. Each data point represents an individual larva, color-coded by replicate. **(C)** The correlation between neutrophil cluster area and fungal area was plotted for larvae that had germination and invasive hyphae. All larvae had neutrophil clusters at some point during infection.

**Table S1.**
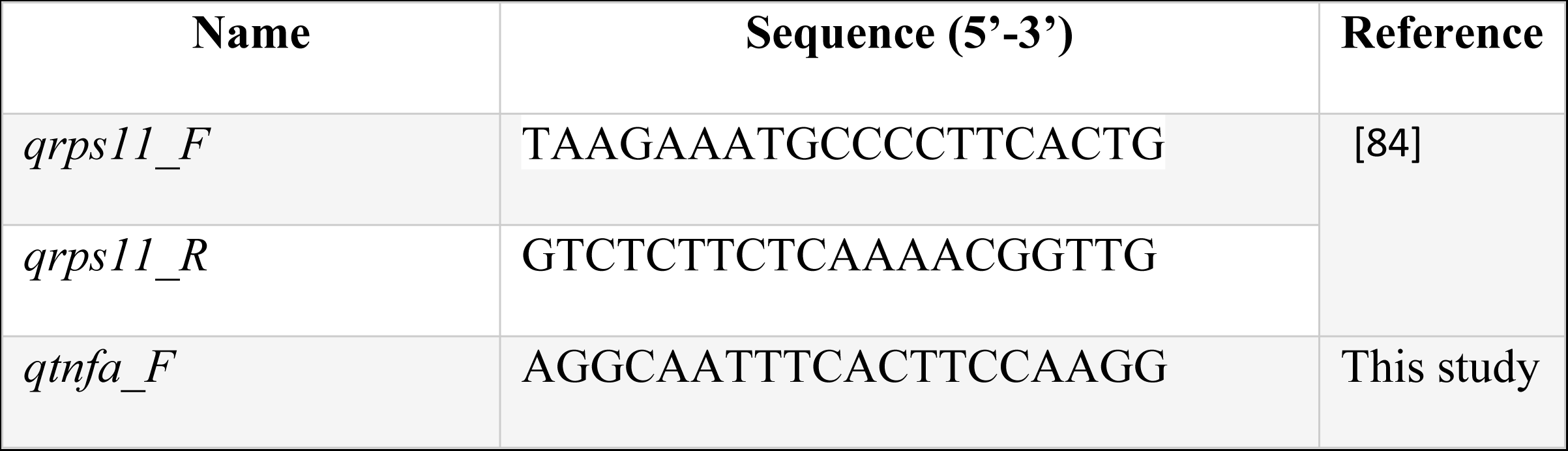
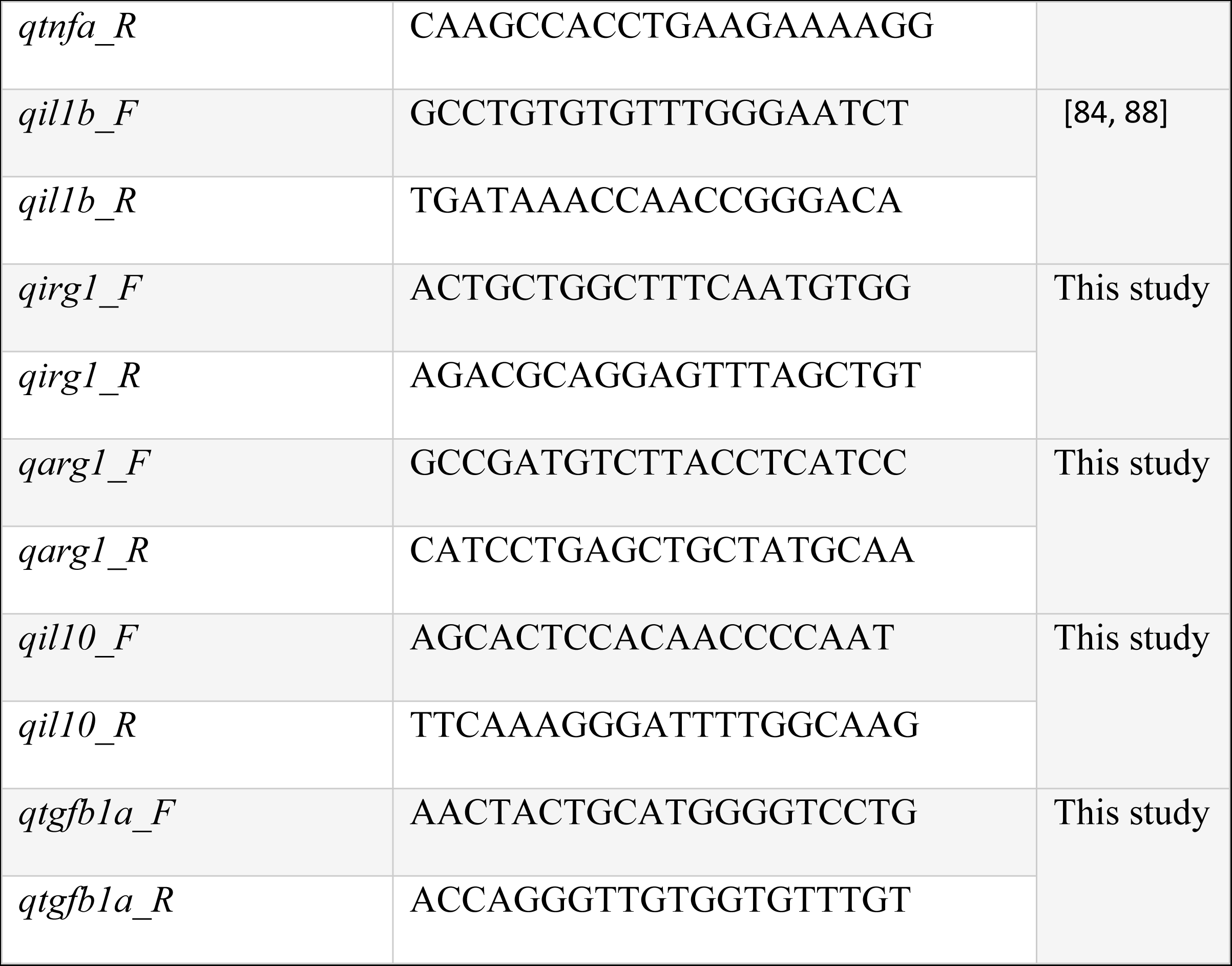
Primers used for RT-qPCR and references.

